# Cells recognize osmotic stress through liquid-liquid phase separation lubricated with poly(ADP-ribose)

**DOI:** 10.1101/2020.04.20.049759

**Authors:** Kengo Watanabe, Kazuhiro Morishita, Xiangyu Zhou, Shigeru Shiizaki, Yasuo Uchiyama, Masato Koike, Isao Naguro, Hidenori Ichijo

**Affiliations:** Laboratory of Cell Signaling, Graduate School of Pharmaceutical Sciences, The University of Tokyo, Tokyo 113-0033, Japan; Department of Cellular and Molecular Neuropathology, Juntendo University Graduate School of Medicine, Tokyo 113-8421, Japan; Department of Cell Biology and Neuroscience, Juntendo University Graduate School of Medicine, Tokyo 113-8421, Japan

## Abstract

Cells are under threat of osmotic perturbation; and cell volume maintenance is critical in cerebral edema, inflammation and aging, in which prominent changes in intracellular or extracellular osmolality emerge. After osmotic stress-enforced cell swelling or shrinkage, the cells regulate intracellular osmolality to recover their volume. However, the mechanisms recognizing osmotic stress remain obscured. We previously clarified that apoptosis signal-regulating kinase 3 (ASK3) bidirectionally responds to osmotic stress and regulates cell volume recovery. Here, we report that macromolecular crowding induces liquid-demixing condensates of ASK3 under hyperosmotic stress, which transduce osmosensing signal into ASK3 inactivation. A genome-wide small interfering RNA (siRNA) screen identified an ASK3 inactivation regulator, nicotinamide phosphoribosyltransferase (NAMPT), related to poly(ADP-ribose) signaling. Furthermore, we clarify that poly(ADP-ribose) keeps ASK3 condensates in the liquid phase and enables ASK3 to become inactivated under hyperosmotic stress. Our findings demonstrate that cells rationally incorporate physicochemical phase separation into their osmosensing systems.

## Introduction

When a difference between intracellular and extracellular osmolality develops, cells inevitably become swollen or shrunken following osmotically driven water flow. The abnormal cellular osmoregulation leads to deteriorated pathophysiological conditions observed in cerebral edema, inflammation, cataracts and aging (*1*–*7*). Basically, homeostasis in cell volume is vital for cellular activities and cells have a defense system against the disastrous osmotic stress; cells immediately excrete or intake ions and small organic solutes after hypoosmotic or hyperosmotic stresses, respectively, and recover their volume within minutes to hours by controlling ion channels and transporters (*8*–*10*). Many electrophysiological and pharmacological studies have contributed to the accumulation of knowledge about the effector molecules in cell volume regulation. In contrast, the mechanisms sensing osmotic stress to induce cell volume recovery remain unclear, especially in mammalian cells. Similar to the osmosensors proposed in bacteria, yeasts and plants (*11*–*13*), mechanical changes in/on the cell membrane have recently drawn attention in mammalian cells; for example, membrane stretching under hypoosmotic stress activates mechanosensitive channels such as the transient receptor potential (TRP) channel V4 (*14, 15*). This mechanism can be illustrated by an easy-to-understand signaling schematic with arrows directed from the extracellular side to the intracellular side. However, osmotic stress perturbs not only the cell membrane but also the intracellular ion strength/concentration and macromolecular crowding (*8, 9*); therefore, the existence of intracellular osmosensors may currently be underestimated.

We previously reported that apoptosis signal-regulating kinase 3 (ASK3; also known as MAP3K15) is phosphorylated and activated under hypoosmotic stress and conversely dephosphorylated and inactivated under hyperosmotic stress (*16*). Furthermore, in addition to the rapid, sensitive and reversible nature, this bidirectional response of ASK3 orchestrates proper cell volume recovery under both hypoosmotic and hyperosmotic stresses (*17*). Therefore, we conceived that the elucidation of ASK3 regulation under osmotic stress would lead to the clarification of a general mammalian osmosensing system. In this study, we report that ASK3 forms liquid droplets under hyperosmotic stress, which is necessary for ASK3 inactivation. Moreover, by utilizing a genome-wide small interfering RNA (siRNA) screen, we reveal that poly(ADP-ribose) (PAR) maintains the liquidity of ASK3 droplets for ASK3 inactivation. Our findings reveal that cells recognize osmotic stress by utilizing liquid-liquid phase separation (LLPS) of ASK3 with the support of PAR.

## Results

### Hyperosmotic stress induces ASK3 condensation through liquid-liquid phase separation

Through analyses of ASK3, we found that the subcellular localization of ASK3 drastically changes under hyperosmotic stress: a part of ASK3 diffuses throughout the cytosol, while the other forms granule-like structures, ASK3 condensates (Fig. 1A). The number of ASK3 condensates increased in a hyperosmolality strength-dependent manner (Fig. 1B) and gradually diminished several dozen minutes after hyperosmotic stress (Fig. S1A and B), which corresponds to the time range of cell volume recovery (*10, 17*). In addition to mannitol-supplemented medium, sodium chloride-supplemented medium induced a similar pattern of ASK3 localization (Fig. S1C and D), suggesting that hyperosmolality causes ASK3 condensates. Counterintuitively, the size of condensates was inversely correlated with hyperosmolality (Fig. 1B). In fact, a simple computational model (*18*) predicted that the size of ASK3 clusters would increase as the grid space is reduced, mimicking cell shrinkage under hyperosmotic stress (Fig. S1E and F, see also Supplementary Text). However, there are abundant macromolecules in cells (*19*); and we modified the model by adding obstacles (Fig. S1G). Our simulation results demonstrated that decreasing the grid space progressively increases both the number and size of ASK3 clusters, while further decreasing the grid space eventually reduces the size of clusters after the maximum has been reached (Fig. 1C, D and Movie S1), implying that the macromolecular crowding is critical for ASK3 condensates under hyperosmotic stress in cells.

**Fig. 1.**
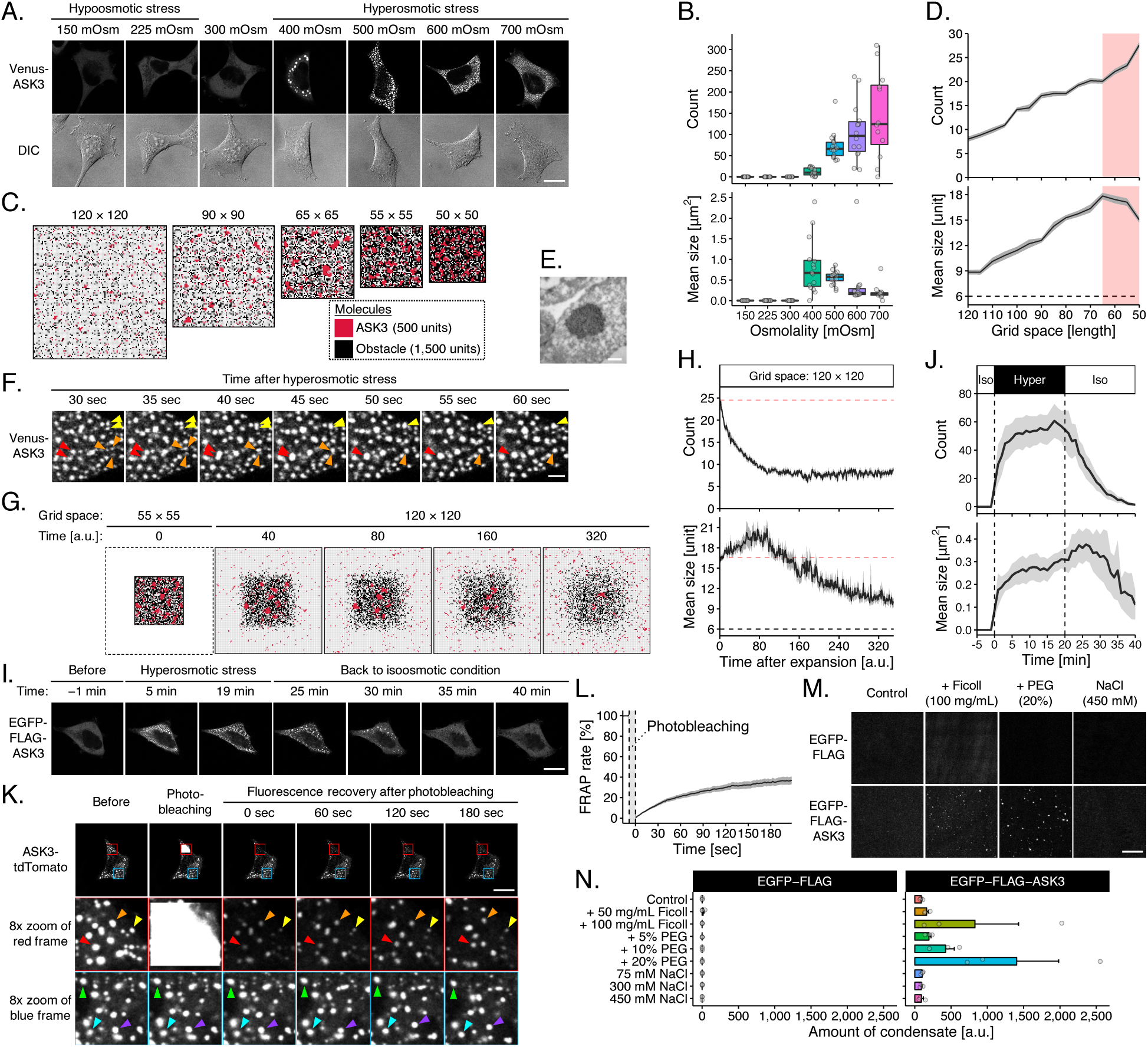
ASK3 forms liquid-demixing condensates under hyperosmotic stress. (**A** and **B**) Subcellular localization of ASK3 5 min after osmotic stress in Venus-ASK3-stably expressing HEK293A (Venus-ASK3-HEK293A) cells. Hypoosmotic stress: ultrapure water-diluted medium, hyperosmotic stress: mannitol-supplemented medium, DIC: differential interference contrast, white bar: 20 μm. *n* = 12–16 cells (pooled from 4 independent experiments). Note that the signal intensity of DIC cannot be compared among the images. (**C** and **D**) A computational simulation for the relationship between the grid space and the number/size of ASK3 clusters. Pale red shading: the assumed range corresponding to hyperosmotic stress (details in Supplementary Text), dashed line: the minimum of clusters definition. Data: mean ± SEM, *n* = 18. (**E**) TEM analysis with immunogold labelling for ASK3. Venus-ASK3-HEK293A cells were sampled after hyperosmotic stress (800 mOsm, 3 hr). White bar: 250 nm. (**F**) Dynamics and fusion of ASK3 condensates in Venus-ASK3-HEK293A cells. Hyperosmotic stress: 500 mOsm, white bar: 2 μm. (**G** and **H**) A computational prediction for the number/size of ASK3 clusters after grid space expansion. Pale red dashed line: the initial values, black dashed line: the minimum of clusters definition. Data: mean ± SEM, *n* = 12. (**I** and **J**) Reversibility of ASK3 condensates in EGFP-FLAG-ASK3-transfected HEK293A cells. After hyperosmotic stress (600 mOsm, 20 min), the extracellular osmolality was set back to the isoosmotic condition. White bar: 20 μm. Data: mean ± SEM, *n* = 8 cells (pooled from 3 independent experiments). (**K** and **L**) FRAP assay for ASK3 condensates in ASK3-tdTomato-transfected HEK293A cells. Prior to the assay, cells were exposed to hyperosmotic stress (600 mOsm, 30 min). White bar: 20 μm. Data: mean ± SEM, *n* = 15 cells (pooled from 5 independent experiments). (**M** and **N**) ASK3 condensation in vitro. Control: 150 mM NaCl, 20 mM Tris (pH 7.5), 1 mM DTT, 15-min incubation on ice. White bar: 10 μm. Data: mean ± SEM, *n* = 3.

Further characterization revealed that ASK3 condensates are colocalized with a marker of neither early endosomes nor lysosomes (Fig. S1H and I). Transmission electron microscopy (TEM) analysis with the immunogold-labelling technique revealed that ASK3 condensates are membrane-less structures (Fig. 1E). Although stress granules and P-bodies are known membrane-less structures under extreme hyperosmotic conditions (*20, 21*), markers of neither structure were found to be colocalized with ASK3 condensates (Fig. S1J). Upon observing in 1-second intervals, we found that ASK3 condensates appear just seconds after hyperosmotic stress, which is much faster than in the case of stress granules, and that ASK3 condensates dynamically move around and fuse with each other (Fig. 1F and Movie S2). Furthermore, our computational model predicted that the shrinkage-induced clusters gradually disappear when the grid space is reverted back to the initial state (Fig. 1G, H and Movie S3, and Supplementary Text), and we observed similar kinetics of reversibility in cells (Fig. 1I and J). Interestingly, our model also predicted a transient increase in the size of clusters just after restoration to the initial space, which was in good agreement with the cell-based experiments. To further address the dynamics of ASK3 condensates, we established a fluorescence recovery after photobleaching (FRAP) assay for the rapidly moving condensates and found that ASK3 condensates display not complete but significant FRAP (Fig. 1K and L), indicating that ASK3 molecules in the condensates are interchanged with those in the cytosol.

Given these characteristics, we concluded that ASK3 condensates are liquid-demixing droplets induced by LLPS (*22*–*24*). Indeed, according to soft matter physics, there are two modes of LLPS, “nucleation and growth” and “spinodal decomposition”, and we observed the spinodal decomposition-like pattern of ASK3 as a rare case (Fig. S1K). Additionally, crowding reagents, such as Ficoll and polyethylene glycol (PEG), induced ASK3 condensates in vitro (Fig. 1M and N). Although the intracellular ion strength and concentration are altered under hyperosmotic stress in cells, the change in sodium chloride concentration did not induce ASK3 condensates in vitro, suggesting again that molecular crowding but not ion strength is a critical driving force for the formation of ASK3 condensates under hyperosmotic stress. Of note, the condensates produced by our in vitro assays are solid-like because we could not observe their FRAP, but the results can be extrapolated to the case in the liquid phase because the driving force in the formation step would be common between the liquid-like and solid-like condensates (*22*–*24*).

### C-terminus coiled-coil and low complexity region are required for ASK3 condensation followed by ASK3 inactivation under hyperosmotic stress

To clarify the significance of ASK3 condensation in cells, we first generated ASK3 mutants that are unable to condense. While ASK3 ΔN normally condensed, ASK3 ΔC lost the ability to condense under hyperosmotic stress (Fig. 2A and B). Moreover, ASK3 CT formed condensates even under isoosmotic conditions, while ASK3 NT and KD did not exhibit the ability to condense, suggesting that the C-terminus of ASK3 is necessary and sufficient for ASK3 condensation. In the ASK3 C-terminus, 5 distinctive regions are bioinformatically predicted (Fig. S2A): two intrinsically disordered regions (IDRs; Fig. S2B), one coiled-coil domain (CC), one sterile alpha motif domain (SAM) (*25*) and one low complexity region (LCR). Between ASK3 CT mutants with these deletions, CTΔCC, CTΔSAM and CTΔLCR exhibited reduced condensation ability compared with the original CT (Fig. S2C). Considering the location of LCR within SAM, we next deleted both CC and SAM or LCR from ASK3 CT and found that CTΔCCΔSAM and CTΔCCΔLCR are unable to condense even under hyperosmotic stress (Fig. S2D), suggesting that both CC and LCR contribute to ASK3 condensation. In addition, we confirmed that full-length ASK3 lacking the C-terminus CC (CCC) and LCR (CLCR) cannot form condensates under hyperosmotic stress (Fig. 2C).

**Fig. 2.**
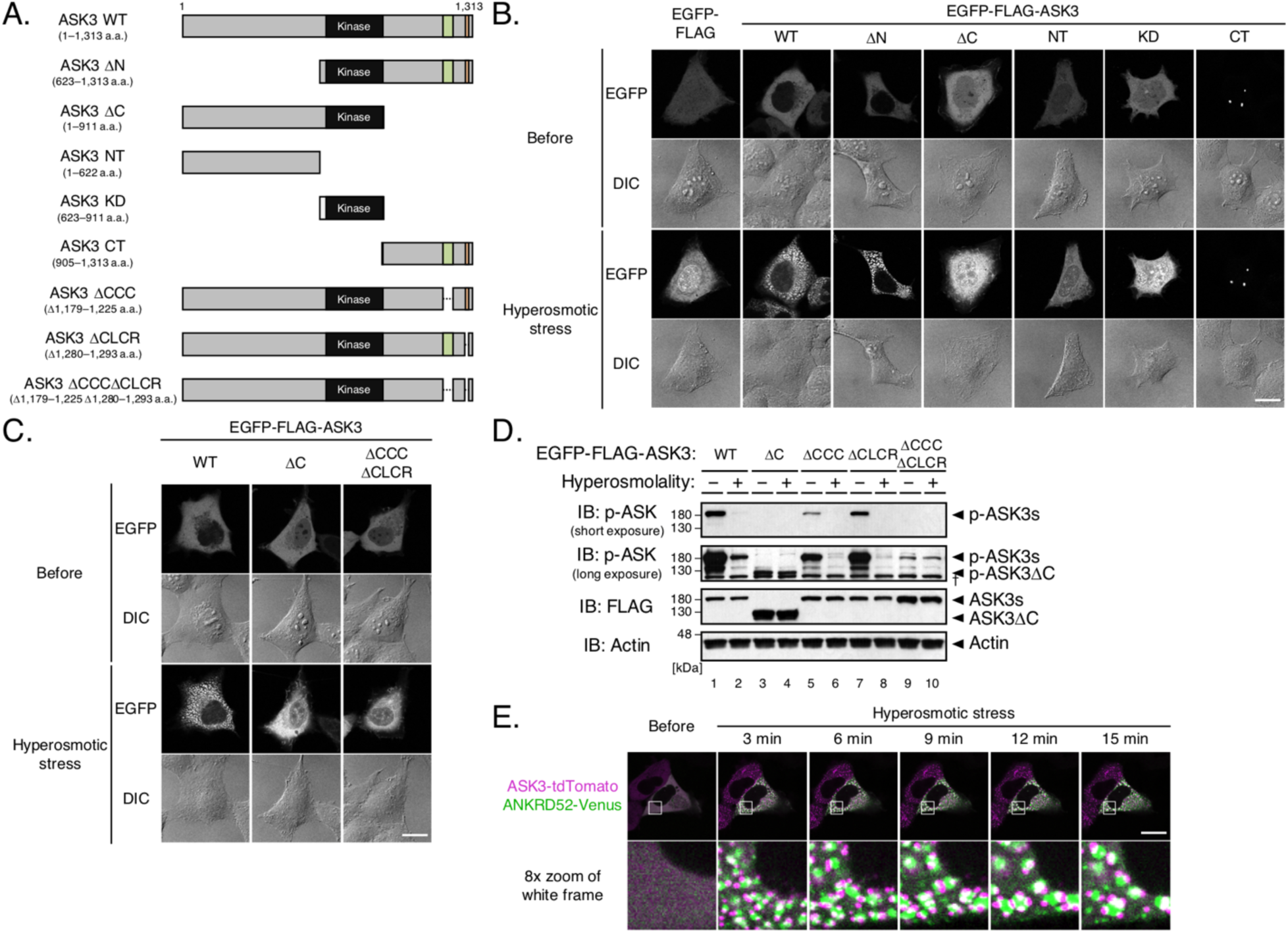
ASK3 condensation is required for ASK3 inactivation under hyperosmotic stress. (**A**) Schematic representation of ASK3 deletion mutants. The numbers indicate the amino acid (a.a.) positions in wild-type (WT). Black rectangle: kinase domain (652–908 a.a.), green rectangle: C-terminus coiled-coil domain (CCC: 1,179–1,225 a.a.), orange rectangle: C-terminus low complexity region (CLCR: 1,280–1,293 a.a.). (**B** and **C**) Subcellular localization of ASK3 mutants in HEK293A cells. Hyperosmotic stress: 600 mOsm, 10 min, DIC: differential interference contrast, white bar: 20 μm. Note that the signal intensity of DIC cannot be compared among the images. (**D**) Requirement of CCC and CLCR for ASK3 inactivation in HEK293A cells. Hyperosmolality (–): 300 mOsm; (+): 500 mOsm; 10 min. IB: immunoblotting. †Nonspecific bands. (**E**) Relationship between ANKRD52 and ASK3 condensates in HEK293A cells. Magenta: ASK3-tdTomato, green: ANKRD52-Venus, hyperosmotic stress: 500 mOsm, white bar: 20 μm.

Previously, we elucidated that ASK3 is inactivated a few minutes after hyperosmotic stress by protein phosphatase 6 (PP6) (*17*). Since ASK3 condensation occurs prior to its inactivation, we evaluated the inactivation of ASK3 mutants lacking condensation ability. Although exhibiting lower basal activities under isoosmotic conditions, ASK3 ΔC and ΔCCCΔCLCR were not inactivated under hyperosmotic stress (Fig. 2D and S3A), suggesting that ASK3 condensation is required for its inactivation. In fact, we discovered the unique relationship between ASK3 condensates and one of the PP6 subunits ANKRD52 (*17, 26*); ANKRD52 condensates are not completely colocalized with ASK3 condensates, but they move around and grow while sharing their phase boundaries (Fig. 2E and Movie S4).

### Nicotinamide phosphoribosyltransferase regulates ASK3 inactivation via the NAD salvage pathway

To reveal the details of ASK3 condensation and inactivation, we investigated the candidate regulators of ASK3 inactivation identified by our genome-wide siRNA screen (*17*). Among them, we focused on the highest-ranked and unexpected candidate nicotinamide phosphoribosyltransferase (NAMPT), the rate-limiting enzyme in the mammalian nicotinamide adenine dinucleotide (NAD) salvage pathway (*27*) (Fig. 3A and B). NAMPT knockdown suppressed ASK3 inactivation under hyperosmotic stress (Fig. 3C and S3B). Additionally, NAMPT knockdown increased endogenous ASK3 activity and inhibited downstream STE20/SPS1-related proline/alanine-rich kinase (SPAK)/oxidative stress-responsive kinase 1 (OSR1) activities under hyperosmotic stress (Fig. 3D and S3C), consistent with our previous finding that ASK3 suppresses SPAK/OSR1 in a kinase activity-dependent manner (*16*). Overexpression of wild-type (WT) NAMPT fully accelerated ASK3 inactivation under hyperosmotic stress in an amount-dependent manner (compare lanes 8–10: Fig. 3E and S3D). In accordance with the fact that NAMPT enzymatically functions as a homodimer (*28*), the homodimer-insufficient mutant S199D could not fully promote ASK3 inactivation, and the homodimer-null mutant S200D could not promote any ASK3 inactivation (*29*) (lanes 11– 14: Fig. 3E and S3D). Furthermore, the NAMPT enzymatic inhibitor FK866 (*30*) inhibited ASK3 inactivation, which was canceled by further supplementation with the NAMPT product nicotinamide mononucleotide (NMN) (Fig. 3F and S3E). Moreover, pretreatment with FK866 and NMN suppressed and promoted the interaction between ASK3 and PP6 under hyperosmotic stress, respectively (*17*) (Fig. 3G, 3H, S3F and S3G). Hence, NAMPT ensures ASK3 inactivation by PP6 via the NAD salvage pathway.

**Fig. 3.**
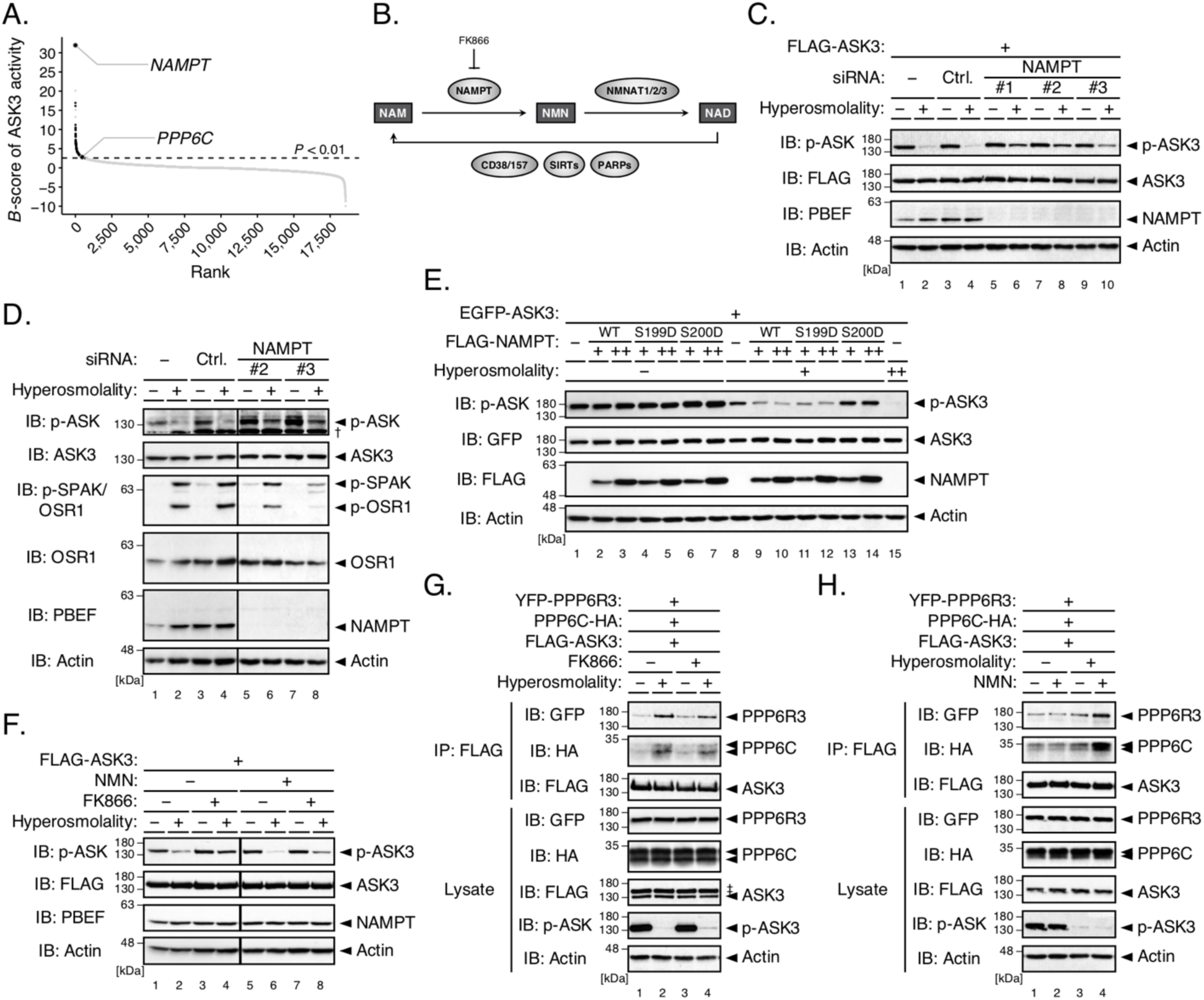
The NAD salvage pathway negatively regulates ASK3 activity under hyperosmotic stress. (**A**) Distribution of gene candidates to regulate ASK3 inactivation in the previous primary screen (*17*). A sample with a higher *B*-score corresponds to a higher potential candidate. PPP6C: the catalytic subunit of PP6, an ASK3 phosphatase. (**B**) Diagram of the NAD salvage pathway. Rectangles, arrows and circles indicate NAD-related molecules, reactions and enzymes, respectively. NAM: nicotinamide, NMN: nicotinamide mononucleotide. (**C**) Effects of NAMPT depletion on ASK3 activity under hyperosmotic stress in FLAG-ASK3-stably expressing HEK293A (FLAG-ASK3-HEK293A) cells. (**D**) Effects of NAMPT depletion on endogenous ASK3 and SPAK/OSR1 activities under hyperosmotic stress in HEK293A cells. †Nonspecific bands. (**E**) Effects of NAMPT overexpression on ASK3 activity under hyperosmotic stress in HEK293A cells. WT: wild-type, S199D: homodimer-insufficient mutant, S200D: homodimer-null mutant. (**F**) Effects of FK866/NMN pretreatment on ASK3 activity under hyperosmotic stress in FLAG-ASK3-HEK293A cells. FK866: 10 nM; NMN: 1 mM; 3-hr pretreatment. (**G** and **H**) Effects of FK866/NMN pretreatment on the interaction between ASK3 and PP6 under hyperosmotic stress in HEK293A cells. FK866: 10 nM; NMN: 1 mM; 24-hr pretreatment. ‡ Remnant bands from prior detection of GFP. (**C**–**H**) Hyperosmolality (–): 300 mOsm; (+): 425 mOsm (with the exception of (G and H), 500 mOsm); (++): 500 mOsm; 10 min, IB: immunoblotting, IP: immunoprecipitation. Note that superfluous lanes were digitally eliminated from blot images in (D and F) as indicated by black lines.

### Poly(ADP-ribose) maintains the liquidity of ASK3 condensates for its inactivation

NAD is not only a coenzyme in cellular redox reactions but also a substrate for enzymatic consumers, including cADP-ribose synthase CD38/157, sirtuins and PAR polymerases (PARPs) (*31*) (Fig. 3B). We thus examined the potential involvement of NAD-consuming enzymes in ASK3 inactivation. In contrast to the overexpression of NAMPT (Fig. 3E), neither CD38, SIRT2 nor PARP1 enhanced ASK3 inactivation under hyperosmotic stress (Fig. S4A–C); rather, their overexpression even suppressed ASK3 inactivation, probably because they compete with the actual NAD-requiring regulators in ASK3 inactivation with respect to NAD. In the dynamics of PARsylation, however, not only PARP writers but also readers and erasers regulate PAR signaling (*32, 33*) (Fig. 4A). Interestingly, NAMPT overexpression and FK866 pretreatment increased and decreased PARsylated proteins, respectively (Fig. 4B and S3H). We therefore examined the potential involvement of PAR signaling in ASK3 inactivation by controlling an eraser PAR glycohydrolase (PARG). PARG overexpression partially suppressed ASK3 inactivation under hyperosmotic stress, while the glycohydrolase-inactive mutant E673A/E674A (*34*) did not (Fig. 4C and S3I), suggesting that PARsylation by unidentified PARP(s) or PAR per se is required for ASK3 inactivation under hyperosmotic stress.

**Fig. 4.**
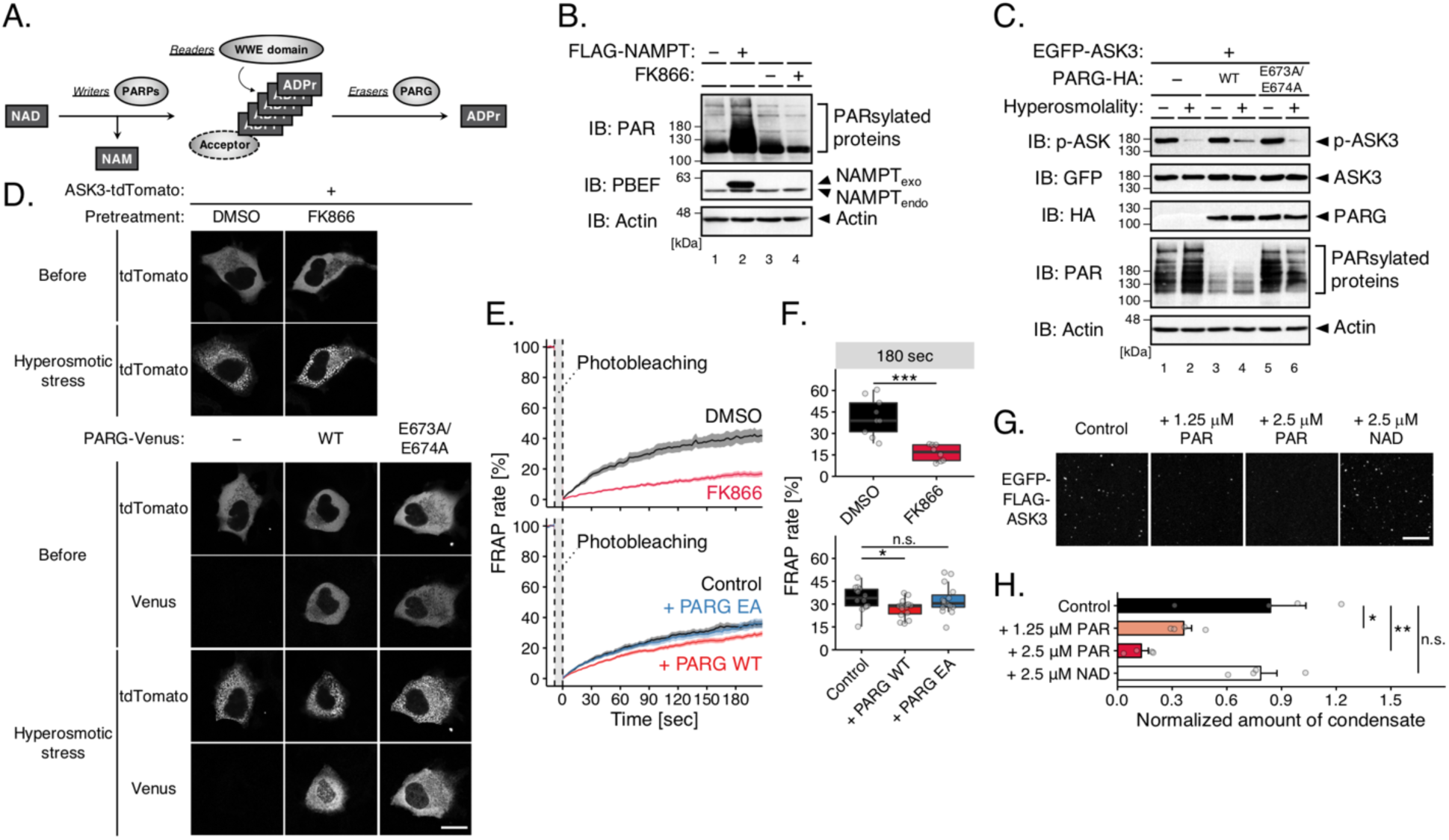
Poly(ADP-ribose) lubricates ASK3 condensates for ASK3 inactivation. (**A**) Diagram of the PARsylation dynamics. Rectangles, arrows and circles indicate NAD-related molecules, reactions and enzymes, respectively. ADPr: ADP-ribose. (**B**) Effects of NAMPT overexpression/inhibition on the amount of PAR in HEK293A cells. FK866: 10 nM, 18–24-hr pretreatment. NAMPT_exo_: exogenously expressed NAMPT, NAMPT_endo_: endogenously expressed NAMPT. (**C**) Effects of PARG overexpression on ASK3 activity under hyperosmotic stress in HEK293A cells. WT: wild-type, E673A/E674A: glycohydrolase-inactive mutant. Hyperosmolality (–): 300 mOsm; (+): 425 mOsm; 10 min. IB: immunoblotting. (**D**) Effects of PAR depletion on ASK3 condensates in ASK3-tdTomato-transfected HEK293A cells. Hyperosmotic stress: 600 mOsm, 10 min, white bar: 20 μm. (**E** and **F**) Effects of PAR depletion on the FRAP of ASK3 condensates in ASK3-tdTomato-transfected HEK293A cells. Prior to the assay, cells were exposed to hyperosmotic stress (600 mOsm, 30 min). Data in (E): mean ± SEM. Top panels: *n* = 8–9 cells (pooled from 3 independent experiments), bottom panels: *n* = 14–15 cells (pooled from 5 independent experiments). \**P* < 0.05, \*\*\**P* < 0.001, n.s. (not significant) by Brunner–Munzel’s tests (with the Bonferroni correction in the bottom). (**D**–**F**) DMSO: negative control for FK866, FK866: 10 nM, 18–24-hr pretreatment, Control: empty vector transfection, PARG WT: wild-type PARG-Venus transfection, PARG EA: E673A/E674A mutant PARG-Venus transfection. (**G** and **H**) Effects of PAR on solid-like ASK3 condensation in vitro. Control: 150 mM NaCl, 20 mM Tris (pH 7.5), 1 mM DTT, 20% PEG, 15-min incubation on ice. White bar: 10 μm. Data: mean ± SEM, *n* = 4. \**P* < 0.05, \*\**P* < 0.01, n.s. (not significant) according to Dunnett’s test.

PAR physicochemically resembles RNA. In addition to RNA, PAR is proposed to seed condensates (*35*–*38*). Considering that ASK3 condensation is required for its inactivation (Fig. 2), we next investigated the relationship between PAR and ASK3 condensation. Contrary to our expectation, ASK3 condensation under hyperosmotic stress was not prevented by PAR depletion with FK866 pretreatment or PARG overexpression (Fig. 4D). However, FRAP of ASK3 condensates was significantly inhibited by PAR depletion (Fig. 4E and F). Furthermore, PAR inhibited the formation of solid-like ASK3 condensates in vitro, while NAD could not (Fig. 4G and H). These results raise the possibility that PAR does not seed but rather lubricates phase-separated ASK3 for ASK3 inactivation.

As a consensus sequence of the PAR-binding motif (PBM), [HKR]_1_-X_2_-X_3_-[AIQVY]_4_-[KR]_5_-[KR]_6_-[AILV]_7_-[FILPV]_8_ is proposed (*39*). Based on the difference between ASK3 WT and CT in condensation at the basal state (Fig. 2B), some kind of inhibitory region is assumed to be present in the region before the C-terminus of ASK3, where the central positively charged [KR]_5_-[KR]_6_ is found in 10 sites, named PBM1–10 (Fig. S5A). We thus constructed 10 PBM candidate mutants substituted K/R with A. All PBM candidate mutants of ASK3 formed condensates under hyperosmotic stress (Fig. S5B). Between them, the ASK3 PBM4 mutant prominently suppressed its inactivation under hyperosmotic stress (Fig. S5C). Indeed, while ASK3 WT was coimmunoprecipitated with PAR by pull-down of the WWE domain (*40*) (Fig. 4A), the PBM4 mutant was not (Fig. S5D), suggesting that ASK3 interacts with PAR via the PBM4 region. Consistent with PAR depletion (Fig. 4E), FRAP of the ASK3 PBM4 mutant was significantly suppressed (Fig. S5E and F). Moreover, the solid-like condensates of the ASK3 PBM4 mutant in vitro were not dissolved by PAR addition (Fig. S5G and H). These findings reinforce the notion that PAR keeps ASK3 condensates in the liquid phase, enabling ASK3 to be inactivated under hyperosmotic stress.

## Discussion

In general, cells face three types of perturbations after osmotic stress: changes in mechanical forces in/on the phospholipid bilayer, intracellular strength/concentration of ions and macromolecular crowding (*9*). It has been suggested that cells recognize unsubstantial osmotic stress through these changes. Here, we demonstrated that an osmoresponsive kinase quickly forms liquid-demixing condensates after hyperosmotic stress, followed by the regulation of its kinase activity. Our computational model and in vitro assays suggested that the change in macromolecular crowding is a driving force for the condensation. Although many researchers have recently been seeking osmosensors in/on the cell membrane, our findings shed light on another mechanism that cells sense osmotic stress from the inside through LLPS. Our findings are reflective of a theory put forth in 1987, when Zimmerman and Harrison proposed “*Changes in reaction rates due to changes in crowding provide, in principle, a simple mechanism by which the cell could sense changes in its own volume*” (*41*). We can advance upon and generalize their idea: cells naturally recognize changes in their volume through phase separation/transition triggered by changes in macromolecular crowding.

Recent studies on LLPS have gradually unveiled the versatile functions of biomolecular condensates, including the acceleration or suppression of specific reactions, the buffering effects on specific biomolecule concentrations, and even the selective filters of nuclear pores (*22, 23, 42*). In this study, we reported the dual function of ASK3 condensates. One function involves the sensing machinery for osmotic stress. Stress-sensing condensates were also suggested in yeast under thermal or pH stresses (*43*). Interestingly, all of these stresses are bidirectionally induced by deviation from steady state; thus, the role of liquid-demixing droplets would be rational in quick and reversible stress recognition. The other is the dephosphorylation/inactivation of ASK3. Because the dephosphorylation competes with the autophosphorylation of ASK3 at the basal level, this would be interpreted as not the trigger of the reaction but the acceleration of the reaction specificity. As another possibility, however, ASK3 condensates may serve as multifunctional signaling hubs for the whole regulation of ASK3 activity. In fact, the ASK3 mutants ΔC and ΔCCCΔCLCR, which are unable to form condensates, exhibited lower kinase activity even under basal conditions (Fig. 2D). Although our confocal microscopy detected few ASK3 condensates under isoosmotic conditions (Fig. 1A), our computational simulation indicated that small clusters appear stochastically and transiently under larger grid space (Fig. 1C and Movie S1), which may be related to a report on local phase separation (*44*). To explore this possibility, further analyses, such as the identification of biomolecules within the ASK3 condensates, are needed.

PAR consists of three components—a base (adenine), ribose and phosphate—which are the same as RNA. In many biomolecular condensates, RNA promotes LLPS of RNA-binding proteins (RBPs) partly because the interaction between RNA and RBPs reduces the threshold for LLPS (*22*–*24*). Likewise, PAR has been reported to promote condensation (*36*–*38*). In contrast, we described that PAR is not a promoter of ASK3 condensates but rather a lubricator. By keeping the liquidity of ASK3 condensates within the “Goldilocks zone”, PAR provides the opportunity for interaction between ASK3 and PP6 followed by ASK3 inactivation; otherwise, ASK3 condensates undergo liquid-to-solid transition (Fig. 4E and S5E), analogous to prion-like RBPs (*45*).

## Materials and Methods

The key materials used in this study are summarized in Table S1.

### Reagents

FK866 (Cat. #F8557) and β-nicotinamide mononucleotide (NMN; Cat. #N3501) were purchased from Sigma-Aldrich and dissolved at a final concentration of 1,000x in dimethyl sulfoxide (DMSO; Sigma-Aldrich, Cat. #D5879) and ultrapure water, respectively. The solvents were used as each negative control.

### Expression plasmids and siRNAs

Expression plasmids for this study were constructed by standard molecular biology techniques, and all constructs were verified by sequencing. Human ASK3 cDNA (CDS of NM_001001671.3 with c.147C>T, c.574G>A) was previously cloned and subcloned into pcDNA3/GW (Invitrogen) with an N-terminal FLAG- or HA-tag (*16*) or into pcDNA4/TO (Invitrogen) with an N-terminal FLAG- or EGFP-tag (*17*). Human ASK3 cDNA was also subcloned into pcDNA3 with a C-terminal tdTomato-tag (cDNA was gifted by M. Davidson, Florida State University, via Addgene: plasmid #54653) or pcDNA4/TO with an N-terminal Venus- or EGFP-FLAG-tag. EGFP-FLAG-tag was constructed by connecting EGFP-tag and FLAG-tag with a Gly-Gly linker and subcloned into pcDNA4/TO. ASK3 mutants ΔN (CDS of NM_001001671.3 with c.1_1,866del), ΔC (CDS of NM_001001671.3 with c.147C>T, c.574G>A, c.2,734_3,939del), NT (CDS of NM_001001671.3 with c.147C>T, c.574G>A, c.1,867_3,939del), KD (CDS of NM_001001671.3 with c.1_1,866del, c.2,734_3,939del), CT (CDS of NM_001001671.3 with c.1_2,712del), ΔCCC (CDS of NM_001001671.3 with c.147C>T, c.574G>A, c.3,535_3,675del), ΔCLCR (CDS of NM_001001671.3 with c.147C>T, c.574G>A, c.3,838_3,879del), ΔCCCΔCLCR (CDS of NM_001001671.3 with c.147C>T, c.574G>A, c.3,535_3,675del, c.3,838_3,879del), CTΔIDR1 (CDS of NM_001001671.3 with c.1_2,712del, c.2,788_2,898del), CTΔIDR2 (CDS of NM_001001671.3 with c.1_2,712del, c.3,427_3,540del), CTΔIDRs (CDS of NM_001001671.3 with c.1_2,712del, c.2,788_2,898del, c.3,427_3,540del), CTΔCC (CDS of NM_001001671.3 with c.1_2,712del, c.3,535_3,675del), CTΔSAM (CDS of NM_001001671.3 with c.1_2,712del, c.3,718_3,909del), CTΔLCR (CDS of NM_001001671.3 with c.1_2,712del, c.3,838_3,879del), CTΔCCΔSAM (CDS of NM_001001671.3 with c.1_2,712del, c.3,535_3,675del, c.3,718_3,909del), CTΔCCΔLCR (CDS of NM_001001671.3 with c.1_2,712del, c.3,535_3,675del, c.3,838_3,879del), PBM1 (R58A/R59A; CDS of NM_001001671.3 with c.147C>T, c.172CGGCGG>GCCGCC, c.574G>A), PBM2 (R203A/R204A; CDS of NM_001001671.3 with c.147C>T, c.574G>A, c.607AGACGA>GCCGCC), PBM3 (R252A/K253A/R255A; CDS of NM_001001671.3 with c.147C>T, c.574G>A, c.754CGGAAA>GCCGCC, c.763AGA>GCC), PBM4 (R332A/R333A; CDS of NM_001001671.3 with c.147C>T, c.574G>A, c.994AGGAGA>GCCGCC), PBM5 (R391A/K392A; CDS of NM_001001671.3 with c.147C>T, c.574G>A, c.1,171CGCAAA>GCCGCC), PBM6 (R424A/K425A; CDS of NM_001001671.3 with c.147C>T, c.574G>A, c.1,270AGGAAA>GCCGCC), PBM7 (R436A/K437A; CDS of NM_001001671.3 with c.147C>T, c.574G>A, c.1,306AGAAAA>GCCGCC), PBM8 (R493A/R494A/K496A/K497A; CDS of NM_001001671.3 with c.147C>T, c.574G>A, c.1,477CGGCG>GCCGC, c.1,686AAGAAA>GCCGCC), PBM9 (K797A/R798A; CDS of NM_001001671.3 with c.147C>T, c.574G>A, c.2,389AAACGT>GCCGCC) and PBM10 (K895A/R896A; CDS of NM_001001671.3 with c.147C>T, c.574G>A, c.2,683AAACGT>GCCGCC) were constructed from full-length ASK3 and subcloned into pcDNA4/TO with an N-terminal EGFP-FLAG-tag. CIDRs in ASK3 were predicted using the IUPred2A tool (*46*) (URL https://iupred2a.elte.hu/) (Fig. S2B). PBMs in ASK3 were defined based on the central positively charged [KR]_5_-[KR]_6_ in the consensus sequence (*38, 39*) (Fig. S5A). The ASK3 PBM4 mutant was also subcloned into pcDNA3/GW with an N-terminal HA-tag or pcDNA3 with a C-terminal tdTomato-tag. The ASK3 CT mutant was also subcloned into pcDNA3 with a C-terminal tdTomato-tag. Human ANKRD52 was cloned previously (*17*) and subcloned into pcDNA3/GW with a C-terminal Venus-tag. Human NAMPT (CDS of NM_005746.2) was cloned from a cDNA pool derived from HEK293A cells and subcloned into pcDNA3/GW with an N-terminal FLAG-tag. NAMPT mutants S199D (CDS of NM_005746.2 with c.595TC>GA) and S200D (CDS of NM_005746.2 with c.598TC>GA) (*29*) were constructed from full-length NAMPT and subcloned into pcDNA3/GW with an N-terminal FLAG-tag. Human PPP6R3 (CDS of NM_001164161.1) with an N-terminal YFP-tag and human PPP6C (CDS of NM_002721.4) with a C-terminal HA-tag were constructed previously (*17*). Human PARG (CDS of NM_001303486.1) and human SIRT2 (CDS of NM_030593.2) were cloned from a cDNA pool derived from HEK293A cells and subcloned into pcDNA3/GW with a C-terminal HA-tag. PARG mutant E673A/E674A (CDS of NM_001303486.1 with c.2,018AAGAA>CCGCC) (*34*) was constructed from full-length PARG and subcloned into pcDNA3/GW with a C-terminal HA-tag. The wild-type PARG and PARG mutant were also subcloned into pcDNA3/GW with a C-terminal Venus-tag. Human PABPC1 (CDS of NM_002568.4) and human DCP1A (CDS of NM_018403.7) were cloned from a cDNA pool derived from HEK293A cells and subcloned into pcDNA3/GW with an N-terminal HA- and Venus-tag, respectively. Human CD38 (CDS of NM_001775.3) was cloned from a cDNA pool derived from A594 cells and subcloned into pcDNA3/GW with a C-terminal HA-tag. Human PARP1 (CDS of NM_001618.3) was cloned from a cDNA pool derived from HeLa cells and subcloned into pcDNA3/GW with an N-terminal HA-tag. WWE domain in human RNF146 (c.247–549 in CDS of NM_030963.3) (*40, 47*) was cloned from a cDNA pool derived from HEK293A cells and subcloned into pcDNA4/TO with an N-terminal EGFP-FLAG-tag. Empty vectors were used as negative controls.

Small interfering RNAs (siRNAs) for human *NAMPT* (#1: 5′-CCACCGACUCCUACAAGGUUACUCA-3′, #2: 5′-GAUCUUCUCCAUACUGUCUUCAAGA-3′, #3: 5′-GAAUAUUGAACUGGAAGCAGCACAU-3′) were purchased as Stealth RNAi siRNAs from Invitrogen. The target sequences were designed using the Block-iT RNAi Designer tool (Invitrogen, current URL https://rnaidesigner.thermofisher.com/rnaiexpress/). As the negative control, Stealth RNAi Negative Control Medium GC Duplex #2 (Invitrogen, Cat. #12935-112) was used.

### Cell culture

HEK293A cells were cultured in Dulbecco’s modified Eagle’s medium (DMEM) (Sigma-Aldrich, Cat. #D5671) supplemented with 10% fetal bovine serum (FBS; BioWest, Cat. #S1560-500) and 100 units/mL penicillin G (Meiji Seika, Cat. #6111400D2039). Tetracycline-inducible Venus-ASK3-stably expressing HEK293A (Venus-ASK3-HEK293A) cells were established with the T-REx system (Invitrogen). Tetracycline-inducible FLAG-ASK3-stably expressing HEK293A (FLAG-ASK3-HEK293A) cells were established previously (*17*). Venus-ASK3-HEK293A cells and FLAG-ASK3-HEK293A cells were cultured in DMEM supplemented with 10% FBS, 2.5 μg/mL blasticidin (Invitrogen, Cat. #A1113903) and 50 μg/mL Zeocin (Invitrogen, Cat. #R25001). To induce Venus-ASK3 or FLAG-ASK3, the cells were pretreated with 1 μg/mL tetracycline (Sigma-Aldrich, Cat. #T7660) 24 hr before assays. All cells were cultured in 5% CO_2_ at 37°C and verified to be negative for mycoplasma.

### Transfections

Plasmid transfections were performed with polyethylenimine “MAX” (Polysciences, Cat. #24765) when HEK293A cells were grown to 95% confluency, according to a previously described protocol (*48*) with minor optimization. To reduce cytotoxicity, the cells were cultured in fresh medium 6–10 hr later, followed by another 40 hr of culture. siRNA transfections for FLAG-ASK3-HEK293A cells were carried out by forward transfection using Lipofectamine RNAiMAX (Invitrogen, Cat. #133778-500) and 10 nM siRNAs once the cells reached 40–80% confluency, according to the manufacturer’s instructions. siRNA transfections for HEK293A cells were carried out by reverse transfection using Lipofectamine RNAiMAX and 30 nM siRNAs, according to the manufacturer’s instructions.

### Osmotic stress treatments

In live-cell imaging experiments, osmotic stress was applied by adding the 2x osmotic medium into the culture medium, followed by the incubation in 5% CO_2_ at 37°C. For isoosmotic conditions (∼300 mOsm/kg H_2_O), DMEM supplemented with 10% FBS was used as the isoosmotic medium. For hyperosmotic stress (∼400, ∼500, ∼600 or ∼700 mOsm/kg H_2_O), DMEM supplemented with 10% FBS and 200, 400, 600 or 800 mM mannitol was used as the 2x hyperosmotic medium. In the case of NaCl-based hyperosmotic stress (∼400, ∼500 or ∼600 mOsm/kg H_2_O), DMEM supplemented with 10% FBS and 100, 200 or 300 mM NaCl was used as the 2x hyperosmotic medium. For hypoosmotic stress (∼150 or 225 mOsm/kg H_2_O), ultrapure water or 2-fold diluted isoosmotic medium was used as the 2x hypoosmotic medium.

In immunoblotting experiments, osmotic stress was applied by exchanging the culture medium with osmotic buffer. The isoosmotic buffer (300 mOsm/kg H_2_O, pH 7.4) contained 130 mM NaCl, 2 mM KCl, 1 mM KH_2_PO_4_, 2 mM CaCl_2_, 2 mM MgCl_2_, 10 mM 4-(2-hydroxyethyl)-1-piperazineethanesulfonic acid (HEPES), 10 mM glucose and 20 mM mannitol. The hyperosmotic buffer (425 or 500 mOsm/kg H_2_O, pH 7.4) was the same as the isoosmotic buffer but contained 145 or 220 mM mannitol, respectively. Absolute osmolality was verified by an Osmomat 030 (Gonotec) osmometer to fall within the range of 295 to 320 mOsm for isoosmotic buffer or ± 25 mOsm/kg H_2_O for the other buffers.

### Live-cell imaging

Cells were seeded in 35 mmφ glass bottom dishes (Matsunami, Cat. #D11130H) coated with 1% gelatin (Nacalai Tesque, Cat. #16605-42) in phosphate-buffered saline (PBS; 137 mM NaCl, 3 mM KCl, 8 mM Na_3_PO_4_•12H_2_O, 15 mM KH_2_PO_4_). For transfected HEK293A cells, the cells were reseeded from a 24-well plate into glass bottom dishes 24 hr after transfection. After 36–60 hr, the culture medium was replaced with 1 mL isoosmotic medium per dish, and the dish was subsequently viewed a TCS SP5 (Leica) confocal laser-scanning microscope equipped with a stage top incubator (Tokai Hit). The cells were observed in 5% CO_2_ at 37°C using an HC PL APO 63x/1.40 oil objective (Leica). Multichannel time-lapse images were acquired in 4 fields each with 4 averages per frame in 1-min or 1.5-min intervals. Venus, EGFP or tdTomato was excited at 514 nm with an argon laser, at 488 nm with an argon laser or at 561 nm with a DPSS laser, respectively, and detected by a HyD detector (Leica). Differential interference contrast (DIC) was captured through the transmitted light from either the argon or DPSS laser, the unused laser in the observation, and detected by a PMT detector (Leica). After obtaining image sets for 5 min under isoosmotic conditions as the “Before” condition, the cells were exposed to osmotic stress by adding 1 mL of 2x osmotic medium per dish and continuously observed for 30 min. Of note, although the cellular morphology was appreciably changed under osmotic stress, we observed the constant position, XY and focal plane, by utilizing a motorized stage and the on-demand mode of adaptive focus control system (Leica) in each field.

In the experiments of the dynamics and fusion of ASK3 condensates, single-channel time-lapse imaging for Venus was performed in a single field with 2 averages per frame at the minimum interval (∼1 sec). To hold a constant position and minimize the autofocusing time, the continuous mode of adaptive focus control system was applied. Similarly, the relationship between ANKRD52 and ASK3 condensates was investigated by 2-channel time-lapse imaging for Venus and tdTomato at the minimum interval (∼5 sec).

In the experiments of the reversibility of ASK3 condensates, time-lapse images of the EGFP and DIC channels were captured by the following procedure. After acquiring the “Before” image sets for 5 min under isoosmotic conditions, the cells were exposed to hyperosmotic stress (600 mOsm) by adding 1 mL of 2x hyperosmotic medium per dish and observed for 20 min. Subsequently, the cells were reverted back to isoosmotic conditions by adding 2 mL of ultrapure water per dish and further observed for 20 min.

For presentation, representative raw images were adjusted in brightness and contrast linearly and equally within the samples by using the GNU Image Manipulation Program (GIMP; GIMP Development Team, URL https://www.gimp.org/) or ImageJ/Fiji (*49*) (URL https://fiji.sc/) software. Because we used DIC images as a rough confirmation of cytosol region, automatically optimized adjustment in brightness and contrast was applied to each DIC image; therefore, the signal intensity of DIC image cannot be compared among the images. To create a time-lapse video, a series of images were equally adjusted in brightness and contrast (and assigned colors if multiple channels were included); captions were added, and the images were converted to a movie file using ImageJ/Fiji software.

For quantification, we established a macro script in ImageJ/Fiji to calculate the count and size of ASK3 condensates in a cell per frame and applied it to all raw image sets in batch mode. Briefly, based on a DIC image, the region of interest (ROI) was first defined as the whole cell area of a main cell because there are condensates from another cell in some cases. After applying a Gaussian filter, the Venus signal within the ROI was subsequently extracted from a Venus image in accordance with the local threshold. Finally, particle analysis was performed. Each parameter was determined from pilot analyses in Venus-ASK3-HEK293A cells. The exported data table was summarized with R language on RStudio (RStudio, Inc., URL https://rstudio.com/) software. The ImageJ/Fiji script also exported images of both ROIs and identified particles, enabling us to confirm the quality. In fact, we excluded several data points from the data analysis: (1) if the image was out-of-focus, (2) if the target cell was shrunken too much or detached completely or (3) if the value was an extreme outlier, less than *Q*_1_ − 5 × IQR or more than *Q*_3_ + 5 × IQR, where *Q*_1_, *Q*_3_ and IQR are the 1st quartile, the 3rd quartile and the interquartile range, respectively.

### Fluorescence recovery after photobleaching assay

Ideally, fluorescence recovery after photobleaching (FRAP) should be applied to only a single condensate. However, ASK3 condensates move around too dynamically and rapidly to be evaluated by normal FRAP assay; ASK3 condensates go out from the focal plane and vice versa, for example. Hence, we established and performed a subsequent FRAP assay for ASK3 condensates under hyperosmotic stress. Prior to the FRAP assay, ASK3-tdTomato-transfected HEK293A cells were placed under the microscope in 5% CO_2_ at 37°C and exposed to hyperosmotic stress (600 mOsm) for 30 min, which makes ASK3 condensates relatively stable. Subsequently, single-channel time-lapse imaging for tdTomato was performed in a single field with 4 averages per frame with a minimum interval (∼2.1 sec) by utilizing the continuous mode of adaptive focus control system. After acquiring 5 frame as the “Before” condition, a rectangular area that included more than 10 condensates (with the exception of the ASK3 CT mutant for which 1 condensate was included) was photobleached by the maximum intensity of the DPSS laser three times, followed by the time-lapse imaging of 100 intervals as the “After” condition.

To quantify the FRAP rate of ASK3 condensates from image data, particle tracking analysis was first executed for all ASK3 condensates by using a Fiji plugin TrackMate (*50*) (URL https://imagej.net/TrackMate). In TrackMate, each condensate was identified in each frame by a Laplacian of Gaussian (LoG) detector, followed by connecting frames by a linear assignment problem (LAP) tracker. Each parameter was determined from pilot analyses for ASK3(WT)-tdTomato. In this tracking analysis, we excluded the condensates (1) that were not successfully tracked from “Before” to “After” or (2) that were present in less than 25 frames. Next, the tracking data table was used to systematically calculated to the FRAP rate in RStudio software. In the R script, each tracked condensate was first categorized into 2 groups, photobleached or not-photobleached, based on the XY coordinates of the photobleached rectangular area. Meanwhile, the fluorescence intensity value of condensate *i* at time *t, F*_*i*_(*t*) was converted to the relative fluorescence change *F*_*i*_(*t*) / *F*_*i*,Before_, where *F*_*i*,Before_ indicates the mean of *F*_*i*_(*t*) for each condensate *i* in the “Before” condition. At this step, we eliminated a few false positive condensates in the photobleached group whose *F*_*i*_(*t*) / *F*_*i*,Before_ did not exhibit at least a 15% decrease between the “Before” and “After” conditions, although *F*_*i*_(*t*) / *F*_*i*,Before_ of the other photobleached condensates dropped by an average of 80% in our FRAP assays. To correct the quenching effects during observation, each *F*_*i*_(*t*) / *F*_*i*,Before_ in the photobleached group was normalized to *G*_*i*_(*t*) = (*F*_*i*_(*t*) / *F*_*i*,Before_) / (the mean of *F*_*j*_(*t*) / *F*_*j*,Before_ in the nonphotobleached group). To mitigate the effects of condensate movement in a direction vertical to the focal plane on the changes in fluorescence, *G*_*i*_(*t*) was further converted to the mean of *G*_*i*_(*t*) in the photobleached group, *G*(*t*); namely, we summarized all values of the photobleached condensates in a cell into representative values of one virtual condensate. Finally, the FRAP rate [%] at time *t* in the cell was calculated as (*G*(*t*) − *G*_Min_) / (1 − *G*_Min_) × 100, where *G*_Min_ was the minimum value within the first three time points of the “After” condition. When summarizing the FRAP rate between cells, we trimmed a few extreme outliers, less than *Q*_1_ − 5 × IQR or more than *Q*_3_ + 5 × IQR.

### Immunocytochemistry and immunofluorescence

Transfected HEK293A cells were seeded on 15 mmφ cover slips (Matsunami, Cat. #C015001) coated with 1% gelatin in PBS in a 12-well plate. After 24–48 hr, the cells were exposed to osmotic medium or buffer for the indicated period, followed by the following immunostaining steps: fixation for 15 min with 4% formaldehyde (Wako Pure Chemical Industries, Cat. #064-00406) in PBS, permeabilization for 15 min with 1% Triton X-100 (Sigma, Cat. #T9284) in PBS, blocking for 30 min with 5% skim milk (Megmilk Snow Brand) in TBS-T (50 mM tris(hydroxymethyl)aminomethane (Tris)-HCl pH 8.0, 150 mM NaCl and 0.05% Tween 20) and incubation at 4°C overnight with the primary antibodies in 1st antibody dilution buffer (TBS-T supplemented with 5% bovine serum albumin (BSA; Iwai Chemicals, Cat. #A001) and 0.1% NaN_3_ (Nacalai Tesque, Cat. #312-33)). The cells were further incubated at room temperature in the dark for 1–2 hr with the appropriate fluorophore-conjugated secondary antibodies in TBS-T. After counterstaining with Hoechst 33258 dye (Dojindo, Cat. #343-07961, 1:2,000) in TBS-T for 5 min, the cover slips were mounted on glass slides with Fluoromount/Plus (Diagnostic Biosystems, Cat. #K048). The samples were observed by using an LSM 510 META (Zeiss) or a TCS SP5 microscope with the 63x/1.40 oil objective. To distinguish the background fluorescence or the “bleed-through” of the other fluorophore from the true signal, we also confirmed the proper negative control samples in each observation.

### Immunoelectron microscopy using ultrathin cryosections

After Venus-ASK3-HEK293A cells were exposed to hyperosmotic stress (800 mOsm, 3 hr), the cells were fixed at room temperature for 10 min with 4% paraformaldehyde (PFA) in 0.1 M phosphate buffer (pH7.2) (PB), followed by the replacement with fresh 4% PFA in PB and incubation at 4°C overnight. Cells were washed 3 times with PBS, followed by 0.15% glycine in PBS, and embedded in 12% gelatin in 0.1 M PB. Small blocks were rotated in 2.3 M sucrose in PB at 4°C overnight and quickly plunged into liquid nitrogen. Sections approximately 60 nm thick were cut using a UC7/FC7 ultramicrotome (Leica) and picked up with a 1:1 mixture of 2% methylcellulose and 2.3 M sucrose in PB. The sections were incubated at 4°C overnight with rabbit anti-GFP antibody (Frontier Institute, Ishikari, Japan, Cat. GFP-Rb-Af2020), followed by incubation at room temperature for 1 hr with protein A conjugated to 10-nm gold particles (protein A-gold; Cell Microscopy Center, University Medical Center Utrecht, Utrecht, the Netherlands). The sections were embedded in a thin layer of 2% methylcellulose with 0.4% uranyl acetate (pH 4.0) and observed with a H-7100 (Hitachi) transmission electron microscope. For control experiments, ultrathin sections were reacted only with protein A-gold.

### In vitro condensation assay

For protein purification, HEK293A cells were seeded in 10 cmφ dishes and transfected with EGFP-FLAG-tagged constructs. After washing with PBS, the cells were lysed in lysis buffer (20 mM Tris-HCl pH 7.5, 150 mM NaCl, 5 mM ethylene glycol-bis(2-aminoethylether)-*N,N,N′,N′*-tetraacetic acid (EGTA), 1% sodium deoxycholate, 1% Triton X-100 and 12 mM β-glycerophosphatase) supplemented with protease inhibitors (1 mM phenylmethylsulfonyl fluoride (PMSF) and 5 μg/mL leupeptin), phosphatase inhibitor cocktail I (8 mM NaF, 1 mM Na_3_VO_4_, 1.2 mM Na_2_MoO_4_, 5 μM cantharidin and 2 mM imidazole) and 1 mM dithiothreitol (DTT). The cell extracts were collected with a scraper from 3 dishes into a single microtube for each protein, followed by centrifugation at 4°C and 13,500 rpm (∼16,500 × *g*) for 15 min. The supernatants were incubated with anti-FLAG antibody beads (Sigma-Aldrich, clone M2, Cat. #A2220) at 4°C for ∼3 hr. The beads were washed 4 times with wash buffer (20 mM Tris-HCl pH 7.5, 500 mM NaCl, 5 mM EGTA, 1% Triton X-100 and 2 mM DTT) and once with TBS (20 mM Tris-HCl pH 7.5, 150 mM NaCl and 1 mM DTT). The EGFP-FLAG-tagged proteins were eluted from the beads with 0.1 mg/mL 3x FLAG peptide (Sigma-Aldrich, Cat. #F4799) in TBS at 4°C for more than 1 hr, followed by dilution to 40 μM with TBS. The concentration of the protein was estimated from the absorbance at 280 nm measured by a SimpliNano (GE healthcare) microvolume spectrophotometer with the extinction coefficient calculated by using the ExPASy ProtParam tool (*51*) (https://web.expasy.org/protparam/).

The purified EGFP-FLAG-tagged protein was diluted into a sample in a microtube, whose control conditions were 10 μM EGFP-FLAG-tagged protein, 150 mM NaCl, 20 mM Tris (pH 7.5), and 1 mM DTT. When increasing the macromolecular crowding, Ficoll PM400 (GE Healthcare, Cat. #17-0310-10) or polyethylene glycol 4000 (PEG; Kanto Kagaku, Cat. #32828-02) was included in the sample at the indicated concentration. When modifying the ion strength and concentration, the concentration of NaCl was changed as indicated. When investigating the effects of poly(ADP-ribose) (PAR) on ASK3 condensates, 20% PEG was added as a crowding reagent, and the indicated concentration of PAR polymer (Trevigen, Cat. #4336-100-01) or β-nicotinamide adenine dinucleotide (NAD; Sigma-Aldrich, Cat. #N7004) in TE (10 mM Tris-HCl pH 8.0 and 1 mM ethylenediaminetetraacetic acid (EDTA)) was also added. The prepared sample was subsequently incubated at 4°C for 15 min. The reaction mixture was immediately loaded into a counting chamber with a cover slip (Matsunami, Cat. # C018241), followed by observation using a TCS SP5 microscope with a 63x/1.40 oil objective. To maintain a constant focal plane even if there were no condensates, we set the focal plane adjacent to the surface of the cover slip by utilizing the motorized stage and the on-demand mode of adaptive focus control system. Images of the EGFP signal were captured from 5 random fields per sample. The above protein purification procedures were performed in each independent experiment. Of note, when we photobleached the fluorescence of ASK3 condensates, we could not observe FRAP; therefore, the condensates produced by our in vitro assays are solid-like, possibly because the maturation of ASK3 condensation occurs too fast in vitro.

For presentation, representative raw images were adjusted in brightness and contrast linearly and equally within the samples using ImageJ/Fiji software. For quantification, we established a macro script in ImageJ/Fiji to calculate the fluorescent intensity and area of ASK3 condensates in each sample and applied the script to all raw image sets in batch mode. In the script, a Gaussian filter and background correction were applied to each image, followed by particle analysis. Each parameter was determined from pilot analyses for EGFP-FLAG-ASK3 WT. The exported data table was further summarized in RStudio software. In the R script, the amount of ASK3 condensates in a sample was defined as the mean of total intensity within 5 fields. When investigating the effects of PAR, the amount value was divided by the amount of the internal standard sample, i.e., the control sample without TE addition. When comparing the effects of PAR between ASK3 mutants, the amount value was converted to the amount relative to the mean of the control sample between experiments.

### Immunoblotting

Cells were lysed in lysis buffer (20 mM Tris-HCl pH 7.5, 150 mM NaCl, 10 mM EDTA, 1% sodium deoxycholate, and 1% Triton X-100) supplemented with protease inhibitors (1 mM PMSF and 5 μg/mL leupeptin). When detecting the phosphorylation of endogenous proteins, phosphatase inhibitor cocktail II (20 mM NaF, 30 mM β-glycerophosphatase, 2.5 mM Na_3_VO_4_, 3 mM Na_2_MoO_4_, 12.5 μM cantharidin and 5 mM imidazole) was also supplemented. When detecting the PARsylated proteins, 1 mM nicotinamide (Sigma-Aldrich, Cat. #N0078) and 100 μM gallotanin (Sigma-Aldrich, Cat. #403040) were also supplemented as PARP and PARG inhibitors, respectively. Cell extracts were clarified by centrifugation at 4°C and 13,500 rpm (∼16,500 × *g*) for 15 min, and the supernatants were sampled by adding 2x sample buffer (80 mM Tris-HCl pH 8.8, 80 μg/mL bromophenol blue, 28.8% glycerol, 4% sodium dodecyl sulfate (SDS) and 10 mM DTT). After boiling at 98°C for 3 min, the samples were resolved by SDS-PAGE and electroblotted onto a BioTrace PVDF (Pall), FluoroTrans W (Pall) or Immobilon-P (Millipore, Cat. #IPVH00010) membrane. The membranes were blocked with 2.5% or 5% skim milk in TBS-T and probed with the appropriate primary antibodies diluted by the 1st antibody dilution buffer. After replacing and probing the appropriate secondary antibodies diluted with skim milk in TBS-T, antibody-antigen complexes were detected on X-ray films (FUJIFILM, Cat. #47410-07523, #47410-26615 or #47410-07595) using an enhanced chemiluminescence system (GE Healthcare). The films were converted to digital images by using a conventional scanner without any adjustment.

For presentation, representative images were acquired by linearly adjusting the brightness and contrast using GIMP software. When digitally trimming superfluous lanes from blot images, the procedure was executed after the adjustment, and the trimming was clearly indicated. Quantification was performed against the raw digital images with densitometry using Fiji/ImageJ software (*17*). “Kinase activity”, “Interaction” and “PARsylated proteins” were defined as the band intensity ratio of phosphorylated protein to total protein, the band intensity ratio of coimmunoprecipitated protein to input protein and the band intensity ratio of PARsylated proteins to actin, respectively. For the kinase activity of endogenous ASK3, the ratio of phosphorylated ASK to ASK3 was calculated.

### Coimmunoprecipitation assay

The supernatants of cell extracts were incubated with anti-FLAG antibody beads (Wako Pure Chemicals Industries, clone 1E6, Cat. #016-22784) for 30–120 min at 4°C. The beads were washed twice with wash buffer (20 mM Tris-HCl pH 7.5, 500 mM NaCl, 1% sodium deoxycholate, 1% Triton X-100, 0.2% SDS) and once with lysis buffer, followed by the direct addition of 2x sample buffer.

### Computational model and simulation

To understand the essential principles of ASK3 condensation in a cell under hyperosmotic stress, we utilized and developed the previously reported simple computational model (*18*) with the effects of macromolecule crowding (Fig. S1G). According to the previous model, each single unit of self-associating ASK3 protein is regarded as an even square that occupies a lattice in a two-dimensional grid space corresponding to a single cell. When an ASK3 unit is adjacent to another, the ASK3 units have a binding relationship, which is independent of other binding pairs. Each ASK3 unit has no relationship between nonadjacent ASK3. A cluster of consecutive multiple ASK3 units is considered as a condensate in cells, which can be assumed to be a liquid-like or solid-like condensate depending on the system parameters mentioned below. At each time step, an ASK3 unit can move to an adjacent lattice. Given the original position of the ASK3 unit, this movement is categorized into three physicochemical actions: (1) diffusion, (2) exchange/vibration and (3) unbinding. If there are no neighboring ASK3 units before the movement (the destination 1 in Fig. S1G), the movement is considered simple diffusion with the rate constant *k*_1_. If there is more than one neighboring ASK3 unit before the movement and if the destination position is already occupied by one of the neighbors (the destination 2 in Fig. S1G), the movement corresponds to an exchange between the ASK3 units in a cluster, namely, a rearrangement process within a liquid-like condensate. This exchange action is obeyed by the rate constant *k*_2_. When *k*_2_ is small, the action is considered vibration of ASK3 in a solid-like condensate. If there is more than one neighboring ASK3 units before the movement and if the destination position is occupied by none of the neighbors (the destination 3a–c in Fig. S1G), the movement can be accompanied by breakages of the binding relationships with the neighbors, i.e., the unbinding reaction. According to the simple Arrhenius equation, a rate constant of this overall unbinding movement *k*_3_ is defined as *k*_3_ = *A* × exp(− Δ*E* × *n*_lost_ / *θ*), where *A* is a frequency factor, Δ*E* is an activation energy in the unbinding reaction between a single pair of ASK3 units, *n*_lost_ is the number of neighboring ASK3 units whose binding relationship with the ASK3 unit to move will be lost by the movement, and *θ* is a temperature-like constant. Notably, the unbinding movement of ASK3 unit includes not only the dissociation process from the condensate but also the rearrangement process of the condensate (compare the destination 3a with the destinations 3b and 3c in Fig. S1G). Moreover, due to the penalty factor *n*_lost_, the rearrangement that increases the surface of condensate (regarded as blue-colored positions in Fig. S1G for the clusters of 2 ASK3 units, for example) is less likely than the rearrangement that maintains in most cases (consider all potential patterns of the case when the destination 4b in Fig. S1G is occupied by not obstacle but ASK3 unit, for example). When *n*_lost_ = 0 (the destination 3c in Fig. S1G), that is, *k*_3_ = *A*, the unbinding movement is considered a shape-modified rearrangement process of the condensate with a void unbinding reaction. In our model, obstacles were further added to reflect on the effects of macromolecular crowding in a cell. Each obstacle basically has the same properties as ASK3 unit; the obstacles takes the same size and shape as ASK3 unit, occupies a grid element and can move to an adjoining position. However, each obstacle has neither a binding relationship with ASK3 units nor one with the other obstacles, that is, each obstacle has neither a positive nor a negative effect on the other molecules. Hence, in addition to the above three physicochemical actions of an ASK3 unit, a new action arises: (4) reflection. If the destination position is occupied by an obstacle (the destination 4a and 4b in Fig. S1G), the ASK3 unit is not able to move to the destination; therefore, the ASK3 unit is reflected by the obstacle and “moves to” the original position according to the rate constant *k*_4_. Simultaneously, the movement of an obstacle is categorized into three physicochemical actions: (5) diffusion, (6) reflection by an ASK3 unit and (7) reflection by an obstacle. If the destination position is unoccupied (the destination 5 in Fig. S1G), the movement is simple diffusion of the obstacle according to the rate constant *k*_5_. If the destination position is already occupied by an ASK3 unit or another obstacle (the destination 6 or 7 in Fig. S1G), the movement corresponds to reflection by the ASK3 unit or the other obstacle, and the obstacle “moves to” the original position depending on the rate constant *k*_6_ or *k*_7_, respectively. We note that the ASK3 unit in our model may not necessarily correspond to a monomer of a single ASK3 peptide in the real world but homo-oligomer(s) of ASK3 or even hetero-oligomer(s). Likewise, the obstacle unit in our model is a virtual molecule that corresponds to the integration of subcellular molecules in the real world, such as macromolecules, small molecules and ions.

To simulate with the model in silico, we computed random trajectories of both ASK3 units and obstacles using the rejection kinetic Monte Carlo (rKMC) method in Python language. Briefly, the Phyton script executed the following algorithm. First, all ASK3 units and obstacles were randomly located in the grid space as an initial condition. Next, a target molecule with a chance to move was randomly picked from the union of the ASK3 unit and obstacle populations, followed by the random selection of a potential destination position of the selected target. As mentioned in the above model description, the designation of the target and destination systematically assigned *k*, the rate constant of the potential movement. At the same time, a random number *r* was acquired from the interval [0, 1). If *k* ≥ *r*, the movement was accepted, and the target was renewed at the position of the destination, although the target was “renewed” at the original position in cases of reflection. Otherwise, if *k* < *r*, the movement was denied, and the target stayed at the original position. After this determination, the iteration step number was incremented by one, and the next iteration step began by randomly selecting the next target molecule.

In the simulation, we regarded *k*_4_, *k*_6_ and *k*_7_ as dummy constants, and we ignored their calculations for the reflection processes because the target molecule remained at its original position in all cases. Since the goal of the computational simulation in this study was not to understand the phase of condensates but to understand the driving force of ASK3 condensation, we also regarded *k*_2_ as a dummy constant and ignored its calculation for the exchange/vibration process. We set *k*_1_ = 1; therefore, free diffusion of ASK3 was always accepted. In contrast, we set *k*_5_ = 0.01; therefore, the diffusion of an obstacle was slower than diffusion of ASK3 unit. This assumption is not unusual because each virtually integrated obstacle is considered the average movement of the constituent molecules, whose movement vectors mostly cancel each other out. For the critical penalty-defining constant *k*_3_, we set *A* = 1 to adjust *k*_3_ = 1 under the condition where *n*_lost_ = 0 and fixed Δ*E* / *θ* = 1 for simplicity; hence, the maximum speed of ASK3 for moving within the condensate was identical to *k*_1_ (= 1), the free diffusion of ASK3 out of condensate. Of note, the sampling distribution of *k*_3_ in our simulation well satisfied exponential decay curve, that is, rejection sampling was confirmed. We prepared 500 ASK3 units with or without 1,500 obstacle units in the grid space. To create the condition under osmotic stress, we changed only the grid space, ranging from 50 × 50 to 120 × 120 squares.

For figure presentation, our Python script saved the coordinates of each molecules, which was rendered using RStudio software. For movie presentation, our Python script also saved representative images at every 10,000 or 100,000 iteration steps, defined as the unit of time *t*. To make a time-lapse video, a series of images with added captions were converted to a movie file using ImageJ/Fiji software. For quantification, our Python script also calculated the count and size of the ASK3 clusters at every iteration step. To mimic confocal microscopy observations, a cluster of condensates was defined as a cluster of ≥6 consecutive ASK3 units. The exported data table was summarized in RStudio software.

### Data analysis and statistical analysis

The data are summarized as the mean ± SEM with the exception of boxplots. No statistical method was utilized to predetermine the sample size. Based on the small sample size and the quality-oriented immunoblotting assays, homoscedasticity was assumed unless *P* < 0.01 in an *F*-test or Levene’s test based on the absolute deviations from the median. Statistical tests, the number of samples and the sample sizes are indicated in Table S2. Statistical tests were performed using R with RStudio software, and *P* < 0.05 was considered statistically significant. In several experiments, a few samples were excluded because they satisfied the criteria clearly outlined in the above sections. The investigators were not blinded to allocation during experiments and outcome assessments. The experiments were not randomized. However, the experiments were performed across different passages of cells, and the cells in the control and treated groups were seeded from the same population of cells.

### Data availability

Almost all data are included in the main text or the supplementary materials. Further information and requests for resources and reagents should be directed to K.W. and H.I.

## Supplementary Materials

- Supplementary Text
- Fig. S1. Characteristics of ASK3 condensates.
- Fig. S2. CCC and CLCR of ASK3 are critical for the ability of ASK3 to condense.
- Fig. S3. Quantification of immunoblotting data in main figures.
- Fig. S4. Major NAD-consuming enzymes suppress ASK3 activity under hyperosmotic stress.
- Fig. S5. ASK3 mutants of PAR-binding motif candidates.
- Table S1. List of key resources.
- Table S2. Summary of statistical analysis.
- Movie S1. A computational simulation for the relationship between the grid space and the number/size of ASK3 clusters.
- Movie S2. Dynamics and fusion of ASK3 condensates in Venus-ASK3-stably expressing HEK293A cells.
- Movie S3. A computational prediction for ASK3 cluster disassembly after the grid space expansion.
- Movie S4. Relationship between ANKRD52 and ASK3 condensates in HEK293A cells.

## Acknowledgments

We thank M. Davidson (Florida State University) for providing tdTomato cDNA via Addgene; all current and former members of the Laboratory of Cell Signaling for valuable materials and fruitful discussions.

## Author contributions

K.W., I.N. and H.I. conceptualized and supervised this project. K.W. designed and performed almost all experiments. K.W. and K.M. established the computational model, and K.M. performed the computational simulation. K.M., X.Z. and S.S. helped with some experiments. Y.U. and M.K. performed the TEM analysis. K.W. visualized and statistically analyzed all the data. K.W., I.N. and H.I. wrote the manuscript.

## References and Notes

1. M. L. McManus, K. B. Churchwell, K. Strange, Regulation of cell volume in health and disease. N. Engl. J. Med. 333, 1260–6 (1995).

2. R. L. Rungta et al., The cellular mechanisms of neuronal swelling underlying cytotoxic edema. Cell. 161, 610–621 (2015).

3. V. Compan et al., Cell volume regulation modulates NLRP3 inflammasome activation. Immunity. 37, 487–500 (2012).

4. J. Jantsch et al., Cutaneous Na+ storage strengthens the antimicrobial barrier function of the skin and boosts macrophage-driven host defense. Cell Metab. 21, 493–501 (2015).

5. L. S. King, D. Kozono, P. Agre, From structure to disease: the evolving tale of aquaporin biology. Nat. Rev. Mol. Cell Biol. 5, 687–698 (2004).

6. L. Hooper, D. Bunn, F. O. Jimoh, S. J. Fairweather-Tait, Water-loss dehydration and aging. Mech. Ageing Dev. 136–137, 50–58 (2014).

7. M. D. Allen, D. A. Springer, M. B. Burg, M. Boehm, N. I. Dmitrieva, Suboptimal hydration remodels metabolism, promotes degenerative diseases, and shortens life. JCI Insight. 4 (2019), doi: 10.1172/jci.insight.130949.

8. F. Lang et al., Functional significance of cell volume regulatory mechanisms. Physiol. Rev. 78, 247–306 (1998).

9. E. K. Hoffmann, I. H. Lambert, S. F. Pedersen, Physiology of cell volume regulation in vertebrates. Physiol. Rev. 89, 193–277 (2009).

10. T. J. Jentsch, VRACs and other ion channels and transporters in the regulation of cell volume and beyond. Nat. Rev. Mol. Cell Biol. 17, 293–307 (2016).

11. J. M. Wood, Osmosensing by bacteria. Sci. STKE. 2006, pe43 (2006).

12. K. Tatebayashi et al., Transmembrane mucins Hkr1 and Msb2 are putative osmosensors in the SHO1 branch of yeast HOG pathway. EMBO J. 26, 3521–33 (2007).

13. F. Yuan et al., OSCA1 mediates osmotic-stress-evoked Ca 2+ increases vital for osmosensing in Arabidopsis. Nature. 514, 367–371 (2014).

14. W. Liedtke et al., Vanilloid receptor-related osmotically activated channel (VR-OAC), a candidate vertebrate osmoreceptor. Cell. 103, 525–35 (2000).

15. R. Strotmann, C. Harteneck, K. Nunnenmacher, G. Schultz, T. D. Plant, OTRPC4, a nonselective cation channel that confers sensitivity to extracellular osmolarity. Nat. Cell Biol. 2, 695–702 (2000).

16. I. Naguro et al., ASK3 responds to osmotic stress and regulates blood pressure by suppressing WNK1-SPAK/OSR1 signaling in the kidney. Nat. Commun. 3, 1285 (2012).

17. K. Watanabe, T. Umeda, K. Niwa, I. Naguro, H. Ichijo, A PP6-ASK3 Module Coordinates the Bidirectional Cell Volume Regulation under Osmotic Stress. Cell Rep. 22, 2809–2817 (2018).

18. E. Dine, A. A. Gil, G. Uribe, C. P. Brangwynne, J. E. Toettcher, Protein Phase Separation Provides Long-Term Memory of Transient Spatial Stimuli. Cell Syst. 6, 655–663.e5 (2018).

19. A. P. Minton, How can biochemical reactions within cells differ from those in test tubes? J. Cell Sci. 119, 2863–2869 (2006).

20. N. L. Kedersha, M. Gupta, W. Li, I. Miller, P. Anderson, RNA-binding proteins TIA-1 and TIAR link the phosphorylation of eIF-2 alpha to the assembly of mammalian stress granules. J. Cell Biol. 147, 1431–42 (1999).

21. J. R. Buchan, R. Parker, Eukaryotic Stress Granules: The Ins and Outs of Translation. Mol. Cell. 36, 932–941 (2009).

22. S. F. Banani, H. O. Lee, A. A. Hyman, M. K. Rosen, Biomolecular condensates: Organizers of cellular biochemistry. Nat. Rev. Mol. Cell Biol. 18, 285–298 (2017).

23. Y. Shin, C. P. Brangwynne, Liquid phase condensation in cell physiology and disease. Science (80-.). 357, eaaf4382 (2017).

24. S. Boeynaems et al., Protein Phase Separation: A New Phase in Cell Biology. Trends Cell Biol. 28, 420–435 (2018).

25. S. J. Trevelyan et al., Structure-based mechanism of preferential complex formation by apoptosis signal–regulating kinases. Sci. Signal. 13, eaay6318 (2020).

26. B. Stefansson, T. Ohama, A. E. Daugherty, D. L. Brautigan, Protein phosphatase 6 regulatory subunits composed of ankyrin repeat domains. Biochemistry. 47, 1442–1451 (2008).

27. J. R. Revollo, A. A. Grimm, S. I. Imai, The NAD biosynthesis pathway mediated by nicotinamide phosphoribosyltransferase regulates Sir2 activity in mammalian cells. J. Biol. Chem. 279, 50754–50763 (2004).

28. T. Wang et al., Structure of Nampt/PBEF/visfatin, a mammalian NAD+ biosynthetic enzyme. Nat. Struct. Mol. Biol. 13, 661–2 (2006).

29. J. R. Revollo et al., Nampt/PBEF/Visfatin regulates insulin secretion in beta cells as a systemic NAD biosynthetic enzyme. Cell Metab. 6, 363–75 (2007).

30. M. Hasmann, I. Schemainda, FK866, a Highly Specific Noncompetitive Inhibitor of Nicotinamide Phosphoribosyltransferase, Represents a Novel Mechanism for Induction of Tumor Cell Apoptosis. Cancer Res. 63, 7436–7442 (2003).

31. S. ichiro Imai, L. Guarente, NAD+ and sirtuins in aging and disease. Trends Cell Biol. 24, 464–471 (2014).

32. W. L. Kraus, PARPs and ADP-Ribosylation: 50 Years … and Counting. Mol. Cell. 58, 902–910 (2015).

33. E. Barkauskaite, G. Jankevicius, I. Ahel, Structures and Mechanisms of Enzymes Employed in the Synthesis and Degradation of PARP-Dependent Protein ADP-Ribosylation. Mol. Cell. 58, 935–946 (2015).

34. N. Le May et al., Poly (ADP-Ribose) Glycohydrolase Regulates Retinoic Acid Receptor-Mediated Gene Expression. Mol. Cell. 48, 785–798 (2012).

35. A. K. L. Leung, Poly(ADP-ribose): An organizer of cellular architecture. J. Cell Biol. 205, 613–619 (2014).

36. A. Patel et al., A Liquid-to-Solid Phase Transition of the ALS Protein FUS Accelerated by Disease Mutation. Cell. 162, 1066–1077 (2015).

37. M. Altmeyer et al., Liquid demixing of intrinsically disordered proteins is seeded by poly(ADP-ribose). Nat. Commun. 6, 8088 (2015).

38. L. McGurk et al., Poly(ADP-Ribose) Prevents Pathological Phase Separation of TDP-43 by Promoting Liquid Demixing and Stress Granule Localization. Mol. Cell. 71, 703–717.e9 (2018).

39. J.-P. Gagné et al., Proteome-wide identification of poly(ADP-ribose) binding proteins and poly(ADP-ribose)-associated protein complexes. Nucleic Acids Res. 36, 6959–76 (2008).

40. Z. Wang et al., Recognition of the iso-ADP-ribose moiety in poly(ADP-ribose) by WWE domains suggests a general mechanism for poly (ADP-ribosyl)ation-dependent ubiquitination. Genes Dev. 26, 235–240 (2012).

41. S. B. Zimmerman, B. Harrison, Macromolecular crowding increases binding of DNA polymerase to DNA: An adaptive effect. Proc. Natl. Acad. Sci. U. S. A. 84, 1871–1875 (1987).

42. H. B. Schmidt, D. Görlich, Transport Selectivity of Nuclear Pores, Phase Separation, and Membraneless Organelles. Trends Biochem. Sci. 41, 46–61 (2016).

43. J. A. Riback et al., Stress-Triggered Phase Separation Is an Adaptive, Evolutionarily Tuned Response. Cell. 168, 1028–1040.e19 (2017).

44. D. Bracha et al., Mapping Local and Global Liquid Phase Behavior in Living Cells Using Photo-Oligomerizable Seeds. Cell, 1467–1480 (2018).

45. S. Maharana et al., RNA buffers the phase separation behavior of prion-like RNA binding proteins. Science. 360, 918–921 (2018).

46. B. Mészáros, G. Erdos, Z. Dosztányi, IUPred2A: context-dependent prediction of protein disorder as a function of redox state and protein binding. Nucleic Acids Res. 46, W329–W337 (2018).

47. P. A. DaRosa et al., Allosteric activation of the RNF146 ubiquitin ligase by a poly(ADP-ribosyl)ation signal. Nature. 517, 223–226 (2014).

48. P. A. Longo, J. M. Kavran, M.-S. Kim, D. J. Leahy, Transient mammalian cell transfection with polyethylenimine (PEI). Methods Enzymol. 529, 227–40 (2013).

49. J. Schindelin et al., Fiji: an open-source platform for biological-image analysis. Nat. Methods. 9, 676–82 (2012).

50. J.-Y. Tinevez et al., TrackMate: An open and extensible platform for single-particle tracking. Methods. 115, 80–90 (2017).

51. E. Gasteiger et al., in The Proteomics Protocols Handbook (Humana Press, Totowa, NJ, 2005; http://link.springer.com/10.1385/1-59259-890-0:571), pp. 571–607.

