## Supplementary Materials for "Cells recognize osmotic stress through liquid-liquid phase separation lubricated with poly(ADP-ribose)"

**This PDF file includes:**

Supplementary Text
Figs. S1 to S5
Tables S1 to S2
Captions for Movies S1 to S4

**Other Supplementary Materials for this manuscript include the following:**

Movies S1 to S4

### Supplementary Text

<u>Technical descriptions of the computational simulation results</u>

In Fig. S1E and S1F, we first utilized the previously reported simple computational model (*18*) to understand the driving force of ASK3 condensation in a cell under hyperosmotic stress. Hyperosmotic stress induces many effects on cells (*1*, *8–10*), but the first substantial trigger must be osmotically driven water efflux, followed by compulsive cell shrinkage. Therefore, we changed the grid space without any changes in the other parameters to mimic the initial cellular condition under hyperosmotic stress. We performed our rKMC with $10^6$ iteration steps at each "cell size" ranging from 50 × 50 to 120 × 120 squares. Our simulation results demonstrated that the decrease in the grid space progressively increased both the count and size of ASK3 clusters within the range of 120 × 120 to 85 × 85 squares. While continuously increasing the size, the further decrease from 85 × 85 to 50 × 50 squares reduced the count from the peak at 85 × 85. This phenomenon is explained by the result that the increase in the cluster size led to a decrease in the number of available free ASK3 units to seed new clusters and the result that the clusters easily fused with each other within the restricted grid space. However, the changing pattern was completely different from that of the cell-based experimental results (compare Fig. S1F with Fig. 1B).

Although it is possible that the model is too simple to explain the phenomena in the real world, one of the biggest differences of the assumption in the model from the cellular condition is the lack of the macromolecular crowding in cells (*19*). In fact, ASK3 has a relatively high molecular mass of 147 kDa, and it is easy to speculate that the effects of the molecular crowding on ASK3 are relatively large in cells. We therefore modified the model by adding obstacles to mimic molecular crowding (Fig. S1G). To maintain simplicity, the properties of obstacles were minimized to hold only an effect of size exclusion (details in Materials and Methods). Since the existence of obstacles prevented ASK3 unit movement and resulted in the slower convergence, we increased the number of iteration steps to $5 \times 10^6$ and executed our rKMC at each "cell size" ranging from 50 × 50 to 120 × 120 squares (Fig. 1C, 1D and Movie S1). Similar to the above results, our simulation results indicated that the decrease in the grid space progressively increased both the number and size of ASK3 clusters, although the range was shifted from 120 × 120 to 65 × 65 squares (Fig. 1D). Nevertheless, while continuously increasing the count, the further decrease from 65 × 65 to 50 × 50 squares reduced the size from the peak at 65 × 65; this pattern of change was the same as that observed in the experimental results (Fig. 1B). Comparing Fig. 1D with Fig. 1B, we interpreted that the grid space from 65 × 65 to 50 × 50 (red shading in Fig. 1D) and the space of approximately 120 × 120 were homologous to the condition under hyperosmotic stress and isoosmotic state in cells, respectively. Indeed, this interpretation may be problematic: for example, (1) whereas the reduction of the grid space ranging from 120 × 120 to 65 × 65 squares gradually increased the count and size of clusters, hyperosmotic stress suddenly increased the count and size of condensates in cells (in other words, there was "dark matter" in the range between 120 × 120 and 65 × 65 squares); and (2) whereas there were small clusters even under isoosmotic conditions in silico, there were no condensates under isoosmotic conditions in cells. However, we should take into account the following three points. First, our model consists of the minimum elements and assumptions; hence, it is not surprising that the details are different. Second, in confocal microscopy observations, we fixed the intensity of lasers and the sensitivity of detectors to acquire the overall appearance of ASK3 in cells from hypoosmotic to hyperosmotic stress. When raising the laser intensity and detector sensitivity, we were able to recognize much smaller condensates under hyperosmotic stress, although the intensity and size of larger condensates became too saturated and overestimated. Likewise, we might be able to detect smaller condensates even under isoosmotic conditions if the noise from ASK3 outside of the condensates could be eliminated. Finally, X-axis is plotted as osmolality in Fig. 1B and as the length of the grid space in Fig. 1D. When assuming a simple situation in which the Boyle–van't Hoff equation can be applied, the changes in cell volume are proportional to the inverse of the changes in osmolality. Hence, the interval of the X-axis variable is not the same between Fig. 1B and Fig. 1D. Altogether, the addition of obstacles enabled the model to properly represent the characteristics of ASK3 condensates in cells, which implies that molecular crowding is a critical driving force for hyperosmotic stress-induced ASK3 condensation in cells, although self-association of ASK3 is fundamental in the potential ability of ASK3 to form condensates.

In Fig. 1G and 1H, by utilizing our computational model, we further performed in silico experiments to predict the changes in ASK3 condensates in cells after hyperosmotic stress is suddenly eliminated. We first iterated rKMC with $5 \times 10^6$ steps at $55 \times 55$ squares of the grid space to grow up ASK3 clusters as shown in Fig. 1C and 1D and to determine the initial positions of ASK3 units and obstacles in subsequent simulations. We next expanded the grid space to $120 \times 120$ squares without any changes in the other parameters, including the molecules' position, and began the simulation with $35 \times 10^6$ iteration steps (Fig. 1G, 1H and Movie S3). The results demonstrated that the number of ASK3 clusters immediately and monotonously decreased as time progressed. Although the size of ASK3 clusters also decreased as time passed, it transiently increased just after the grid space was expanded to $120 \times 120$ squares. This interesting phenomenon can be interpreted as follows: obstacles gradually emerge from the original $55 \times 55$ squares after the grid space is extended, accompanied by the enormous decrease in the effects of the size exclusion on ASK3 units. Free ASK3 units can therefore access the surface of clusters, which cause the clusters to grow. At the same time, ASK3 units can also emerge from the original grid space, which permits the dissociation of constitutive ASK3 units from the clusters and decreases the size of the clusters. Hence, these two opposing behaviors compete with each other. Due to the proximity of the ASK3 clusters at the initial phase and the existence of the unbinding penalty, this interesting transition of power balance is observed.

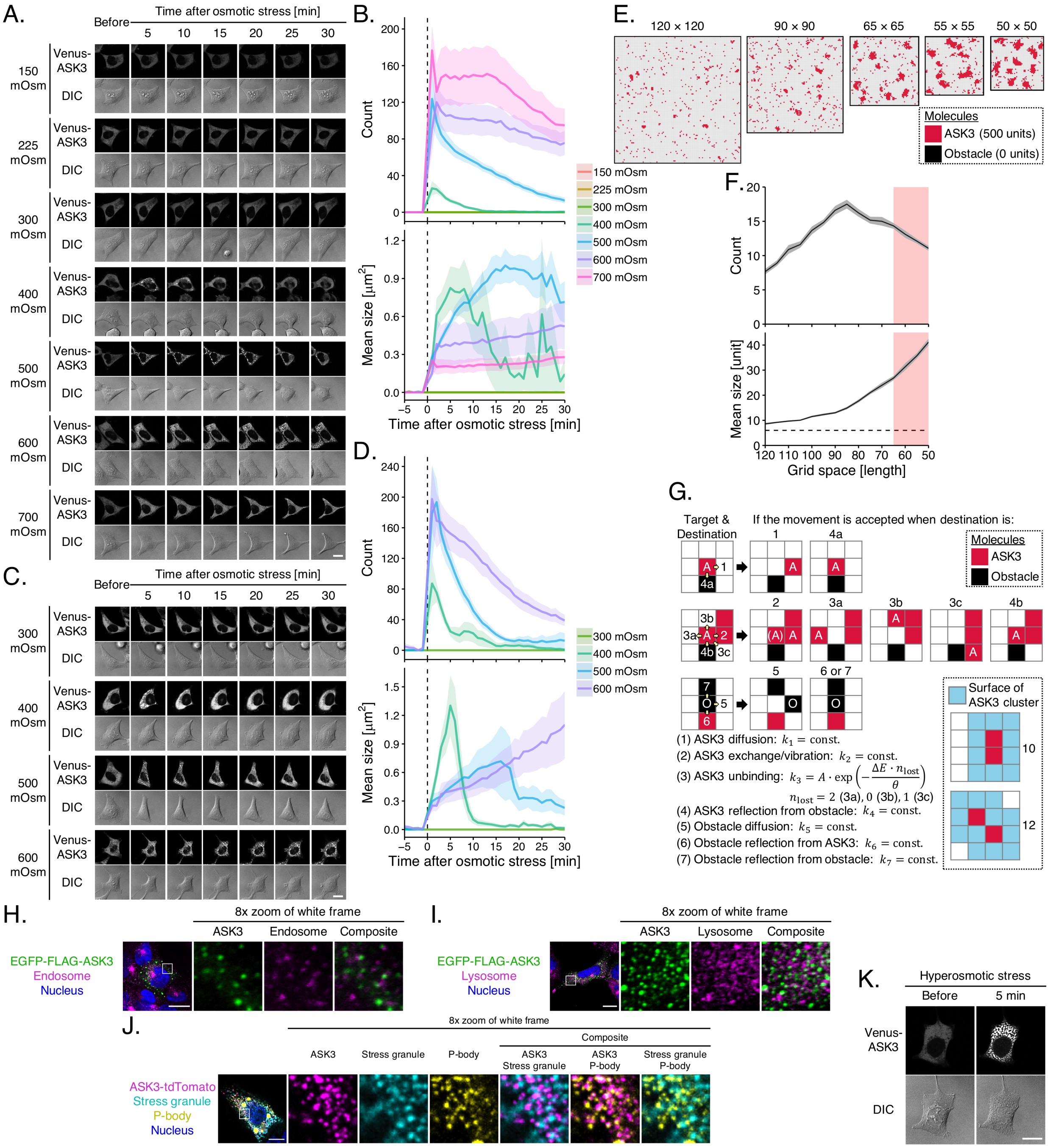


**Fig. S1. Characteristics of ASK3 condensates.**

(**A** and **B**) Time course of the changes in the subcellular localization of ASK3 after osmotic stress in Venus-ASK3-stably expressing HEK293A (Venus-ASK3-HEK293A) cells. Hypoosmotic stress: ultrapure water-diluted medium, hyperosmotic stress: mannitol-supplemented medium, DIC: differential interference contrast, white bar: 20 µm. Data: mean ± SEM, *n* = 12–16 cells (pooled from 4 independent experiments). (**C** and **D**) Effects of sodium chloride on the subcellular localization of ASK3 in Venus-ASK3-HEK293A cells. Hyperosmotic stress: NaCl-supplemented medium, white bar: 20 µm. Data: mean ± SEM, *n* = 10–12 cells (pooled from 3 independent experiments). (**E** and **F**) A computational simulation of the relationship between the grid space and the number/size of ASK3 clusters using the previously reported model (*18*). Pale red shading: corresponding range in Fig. 1D, dashed line: the minimum of clusters definition. (**G**) Schematic diagram of our model. Red squares: ASK3 units, black squares: obstacles, white arrows: potential movements, blue squares: surface positions of the clusters. See Materials and Methods for details. (**H** and **I**) Relationship between ASK3 condensates and early endosomes (H) or lysosomes (I). EGFP-FLAG-ASK3-transfected HEK293A cells were sampled after hyperosmotic stress (500 mOsm, 15 min). Early endosomes: immunofluorescence with an antibody against EEA1, lysosomes: immunofluorescence with an antibody against LAMP1, white bar: 15 µm. (**J**) Relationship between ASK3 condensates and stress granules/P-bodies. Transfected HEK293A cells were sampled after hyperosmotic stress (800 mOsm, 45 min). ASK3: ASK3-tdTomato, stress granules: HA-PABPC1 (immunofluorescence with an antibody against HA-tag), P-bodies: Venus-DCP1A, white bar: 10 µm. (**K**) An example of ASK3 condensates in spinodal decomposition-like pattern. Venus-ASK3-HEK293A cells were exposed to hyperosmotic stress (600 mOsm), white bar: 20 µm. Note that the DIC signal intensity cannot be compared among the images in (A, C and K).

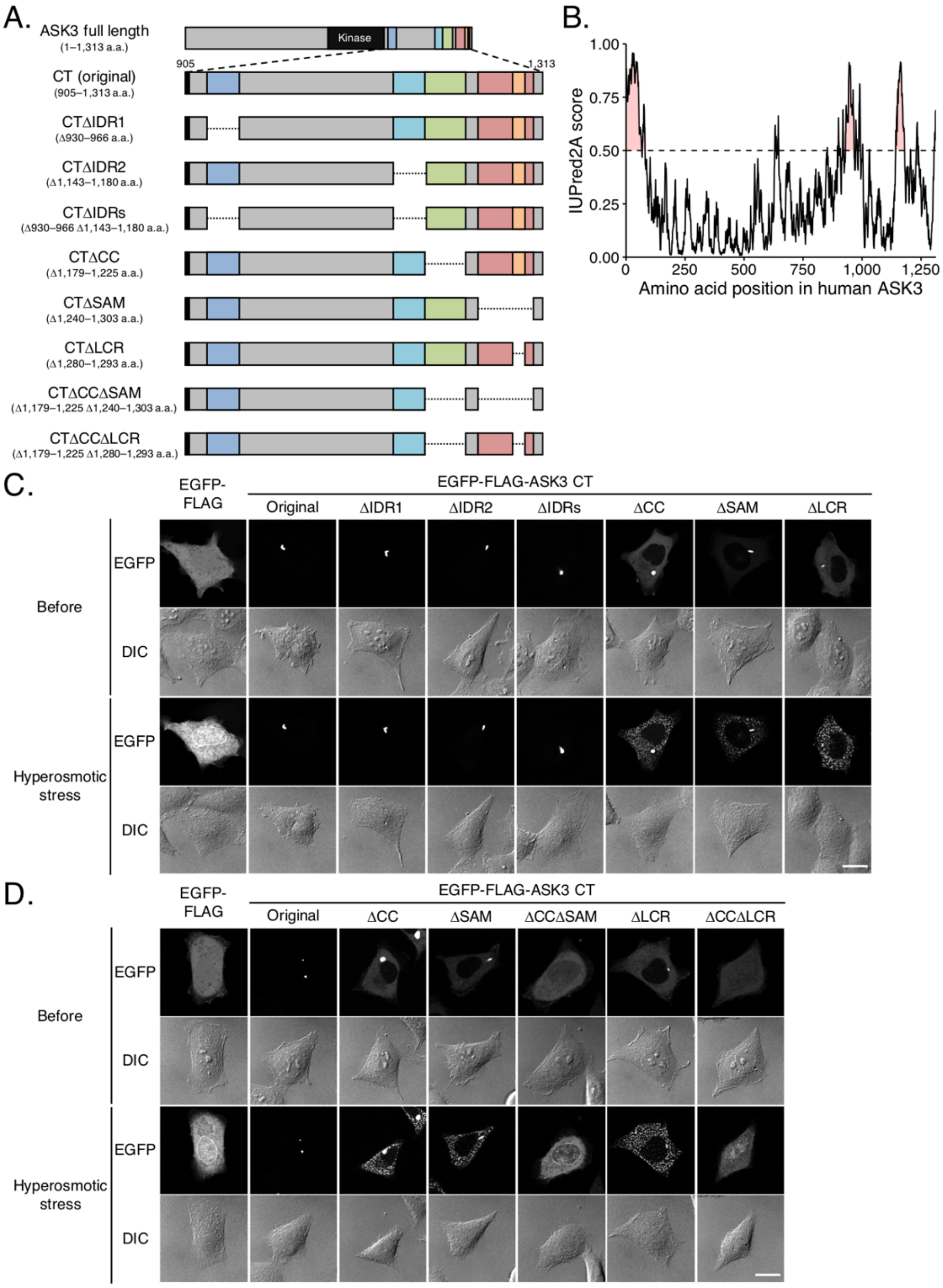


**Fig. S2. CCC and CLCR of ASK3 are critical for the ability of ASK3 to condense.**
(**A**) Schematic representation of ASK3 deletion mutants. The numbers indicate amino acid (a.a.) positions in the full-length wild-type ASK3. Black rectangle: kinase domain (652–908 a.a.), dark blue rectangle: C-terminus intrinsically disordered regions (CIDR1: 930–966 a.a.), light blue rectangle: CIDR2 (1,143–1,180 a.a.), green rectangle: C-terminus coiled-coil domain (CCC: 1,179–1,225 a.a.), red rectangle: sterile alpha motif domain (SAM: 1,240–1,303 a.a.), orange rectangle: C-terminus low complexity region (CLCR: 1,280–1,293 a.a.). (**B**) Prediction of IDRs in ASK3 using the IUPred2A tool (*46*). (**C** and **D**) Subcellular localization of ASK3 CT mutants in HEK293A cells. Hyperosmotic stress: 500 mOsm, 10 min, DIC: differential interference

contrast, white bar: 20 μm. Note that the DIC signal intensity cannot be compared among the images.

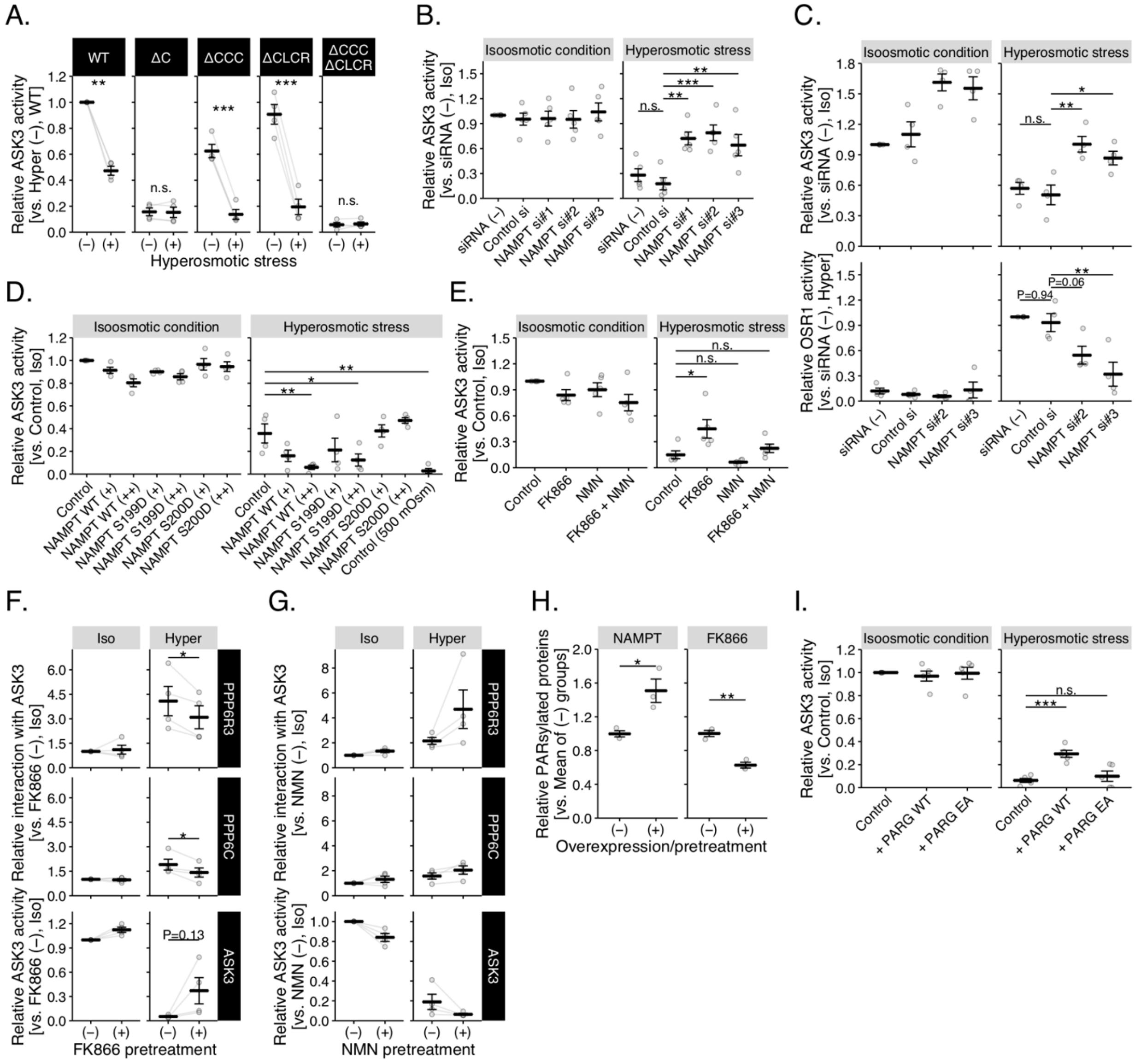


**Fig. S3. Quantification of immunoblotting data in main figures**

(**A**) Quantification of Fig. 2D. (**B**) Quantification of Fig. 3C. (**C**) Quantification of Fig. 3D. (**D**) Quantification of Fig. 3E. (**E**) Quantification of Fig. 3F. (**F**) Quantification of Fig. 3G. (**G**) Quantification of Fig. 3H. (**H**) Quantification of Fig. 4B. (**I**) Quantification of Fig. 4C. Individual values and the mean ± SEM are presented as gray points (connected with gray lines within the same experimental sets in (A, F and G)) and black lines, respectively. **P* < 0.05, ***P* < 0.01, ****P* < 0.001, n.s. (not significant). Statistical tests, the number of samples and the sample sizes are summarized in Table S2.

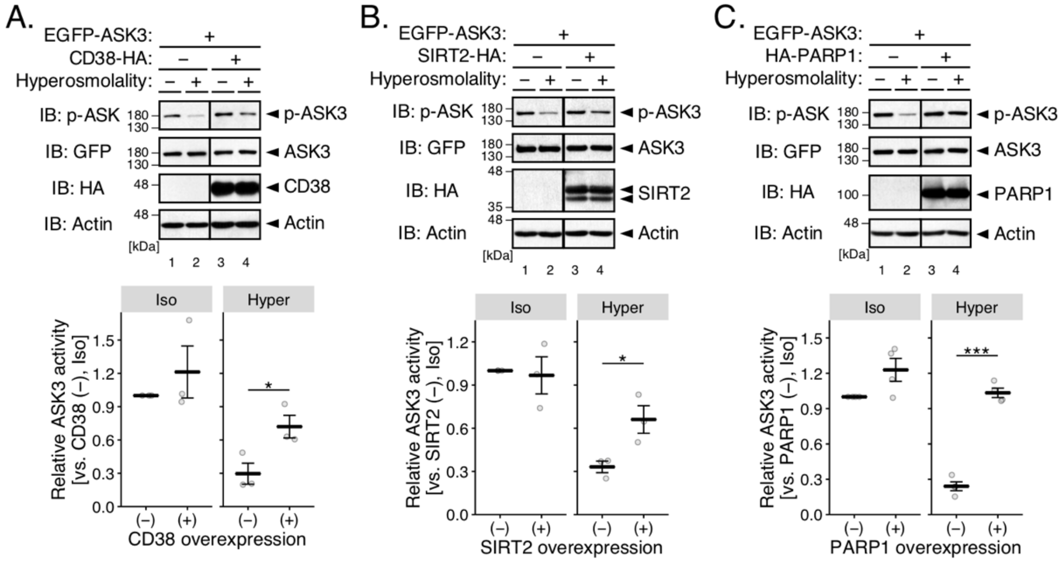


**Fig. S4. Major NAD-consuming enzymes suppress ASK3 activity under hyperosmotic stress.**

(**A**–**C**) Effects of CD38 (A), SIRT2 (B) or PARP1 (C) overexpression on ASK3 activity under hyperosmotic stress in HEK293A cells. The top panel is a representative image set of immunoblotting, and the bottom graph depicts the quantification. Hyperosmolality (–): 300 mOsm; (+): 425 mOsm; 10 min. IB: immunoblotting. Note that superfluous lanes were digitally eliminated from blot images as indicated by black lines. In the bottom graphs, individual values and the mean ± SEM are presented as gray points and black lines, respectively. $*P < 0.05$, $***P < 0.001$. Statistical tests, the number of samples and the sample sizes are summarized in Table S2.

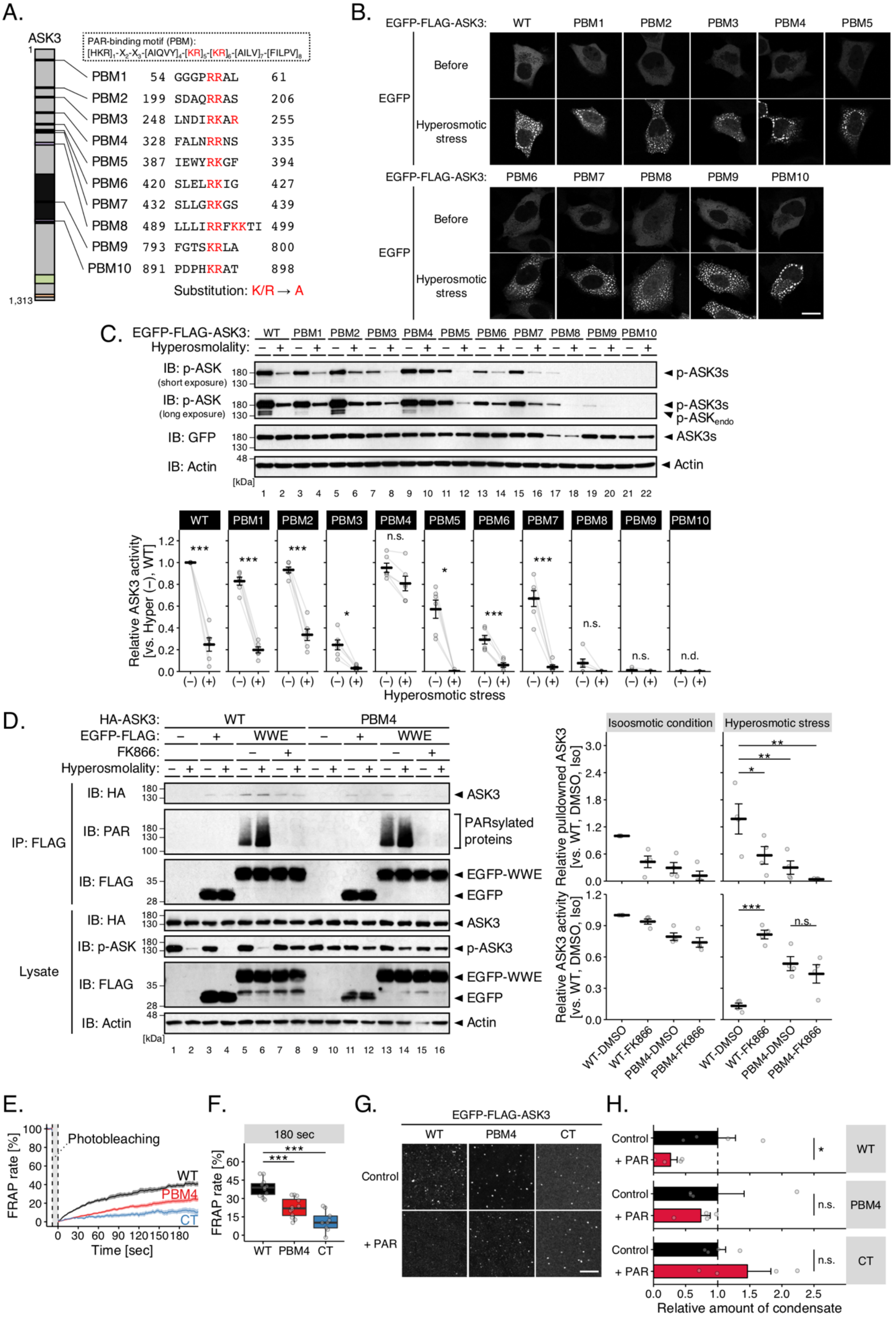
A.
ASK3
PAR-binding motif (PBM):
PBM1 54 GGGPRRAL 61
PBM2 199 SDAQRRAS 206
PBM3 248 LNDIRKAR 255
PBM4 328 FALNRRNS 335
PBM5 387 IEWYRKGF 394
PBM6 420 SLELRKIG 427
PBM7 432 SLLGRKGS 439
PBM8 489 LLLIRRFKKTI 499
PBM9 793 FGTSKRLA 800
PBM10 891 PDPHKRAT 898
Substitution: K/R → A
1,313
B.
EGFP-FLAG-ASK3: WT PBM1 PBM2 PBM3 PBM4 PBM5
EGFP
Before
Hyperosmotic stress
EGFP-FLAG-ASK3: PBM6 PBM7 PBM8 PBM9 PBM10
C.
EGFP-FLAG-ASK3: WT PBM1 PBM2 PBM3 PBM4 PBM5 PBM6 PBM7 PBM8 PBM9 PBM10
Hyperosmolality:
IB: p-ASK (short exposure)
IB: p-ASK (long exposure)
IB: GFP
IB: Actin
◄ p-ASK3s
◄ ASK3s
◄ Actin
[kDa]
Relative ASK3 activity [vs. Hyper (–), WT]
Hyperosmotic stress
n.s.
n.d.
D.
HA-ASK3: WT PBM4
EGFP-FLAG: – + WWE
FK866:
Hyperosmolality:
IP: FLAG
IB: HA
IB: PAR
IB: FLAG
Lysate
IB: p-ASK
IB: Actin
◄ ASK3
PARsylated proteins
◄ EGFP-WWE
◄ EGFP
◄ p-ASK3
◄ Actin
Isoosmotic condition
Hyperosmotic stress
Relative pulldowned ASK3 [vs. WT, DMSO, Iso]
Relative ASK3 activity [vs. WT, DMSO, Iso]
WT-DMSO
WT-FK866
PBM4-DMSO
PBM4-FK866
E.
Photobleaching
FRAP rate [%]
Time [sec]
WT
PBM4
CT
F.
180 sec
FRAP rate [%]
WT PBM4 CT
G.
EGFP-FLAG-ASK3
WT PBM4 CT
Control
+ PAR
H.
Control
+ PAR
WT
PBM4
CT
n.s.
Relative amount of condensate

165

**Fig. S5. ASK3 mutants of PAR-binding motif candidates**

(**A**) Candidate PAR-binding motif (PBM) in ASK3. Schematic representation of ASK3 is the same as that in Fig. 2A. The numbers indicate the amino acid (a.a.) positions in wild-type (WT). (**B**) Subcellular localization of ASK3 PBM candidate mutants in HEK293A cells. Hyperosmotic stress: 500 mOsm, 10 min, white bar: 20 µm. (**C**) Inactivation of ASK3 PBM candidate mutants under hyperosmotic stress in HEK293A cells. p-$ASK_{endo}$: autophosphorylated endogenous ASK3 by exogenously expressed ASK3. (**D**) The ability of the ASK3 PBM4 mutant to interact with PAR under hyperosmotic stress in HEK293A cells. FK866: 10 nM, 12-hr pretreatment. (**C** and **D**) The top or left panel is a representative image set of immunoblotting, and the bottom or right graph depicts the quantification. Hyperosmolality (–): 300 mOsm; (+): 500 mOsm; 10 min. IB: immunoblotting, IP: immunoprecipitation. In the graphs, individual values and the mean ± SEM are presented as gray points (connected with gray lines within the same experimental sets in (C)) and black lines, respectively. $*P < 0.05$, $**P < 0.01$, $***P < 0.001$, n.s. (not significant), n.d. (not detected). Statistical tests, the number of samples and the sample sizes are summarized in Table S2. (**E** and **F**) FRAP assay for the condensates of the ASK3 PBM4 mutant in HEK293A cells. Prior to the assay, cells transfected with each ASK3-tdTomato were exposed to hyperosmotic stress (600 mOsm, 30 min). Data in (E): mean ± SEM. $n = 9–12$ cells (pooled from 4 independent experiments). $***P < 0.001$ by Brunner–Munzel's tests with the Bonferroni correction. (**G** and **H**) Effects of PAR on the solid-like condensates of the ASK3 PBM4 mutant in vitro. Control: 150 mM NaCl, 20 mM Tris (pH 7.5), 1 mM DTT, 20% PEG, 15-min incubation on ice. White bar: 10 µm. Data: mean ± SEM, $n = 4$. $*P < 0.05$, n.s. (not significant) by unpaired two-sample *t*-test.

**Table S1. List of key resources**

Key resources used in this study are indicated. IB: immunoblotting, IF: immunofluorescence, IEM: immunoelectron microscopy, CDS: cording sequence.

| REAGENT or RESOURCE | SOURCE or REFERENCE |
|---|---|
| Antibodies | |
| Rabbit polyclonal anti-phospho-ASK (p-ASK; Thr808 in human ASK3, IB: 1:20,000 or 1:2,000) | (Naguro et al., 2012) |
| Anti-rabbit IgG, HRP-linked (IB: 1:1,000–1:10,000) | Cell Signaling Technology (Cat. #7074) |
| Mouse monoclonal anti-DYKDDDDK tag (FLAG; clone 1E6, IB: 1:5,000 or 1:1,000) | Wako Pure Chemical Industries (Cat. #012-22384) |
| Anti-mouse IgG, HRP-linked (IB: 1:2,000–1:20,000) | Cell Signaling Technology (Cat. #7076) |
| Mouse monoclonal anti-Actin (Actin; clone AC-40, IB: 1:2,000) | Sigma-Aldrich (Cat. #A3853) |
| Rabbit polyclonal anti-PBEF (PBEF; IB: 1:10,000) | Bethyl Laboratories (Cat. #A300-372A) |
| Rat monoclonal anti-ASK3 (ASK3; IB: 10,000) | (Naguro et al., 2012) |
| Anti-rat IgG, HRP-linked (IB: 1:2,000–1:20,000) | Cell Signaling Technology (Cat. #7077) |
| Rabbit polyclonal anti-phospho-SPAK/OSR1 (p-SPAK/OSR1; Thr231 in human SPAK and Thr185 in human OSR1, IB: 1:1,000) | (Naguro et al., 2012) |
| Mouse monoclonal anti-OXSR1 (OSR1; clone 2A2-1A2, IB: 1:20,000) | Abnova (Cat. #H00009943-M01) |
| Mouse monoclonal anti-GFP (GFP; clone 1E4, IB: 1:10,000 or 1:5,000) | Medical & Biological Laboratories (Cat. #M048-3) |
| Rat monoclonal anti-YPYDVPDYA tag (HA; clone 3F10, IB: 1:5,000 or 1:1,000) | Roche Diagnostics (Cat. #11867431001) |
| Rabbit polyclonal anti-Poly(ADP-ribose) (PAR; IB: 1:5,000) | Enzo Life Sciences (Cat. #ALX-210-890A) |
| Anti-DYKDDDDK tag Antibody Beads (FLAG; clone 1E6, IP) | Wako Pure Chemical Industries (Cat. #016-22784) |
| ANTI-FLAG M2 Affinity Gel (clone M2, IP) | Sigma-Aldrich (Cat. #A2220) |
| Mouse monoclonal anti-EEA1 (EEA1; clone 14/EEA1, IF: 1:200) | BD Transduction Laboratories (Cat. #610456) |
| Alexa Fluor 555 goat anti-mouse IgG (H+L) (IF: 1:300) | Molecular Probe (Cat. #A21422) |
| Mouse monoclonal anti-LAMP1 (LAMP1; clone H4A3, IF: 1:100) | Santa Cruz Biotechnology (Cat. #sc-20011) |
| Alexa Fluor 633 goat anti-rat IgG (H+L) (IF: 1:300) | Molecular Probe (Cat. #A21094) |
| Rabbit polyclonal anti-GFP antibody (IEM) | Frontier Institute (Cat. #GFP-Rb-Af2020) |
| Chemicals and Peptides | |
| Polyethylenimine “MAX” | Polysciences (Cat. #24765) |
| Lipofectamine RNAiMAX | Invitrogen (Cat. #13778-500) |
| Zeocin | Invitrogen (Cat. #R25001) |
| Blasticidin S HCl | Invitrogen (Cat. #A1113903) |
| Tetracycline | Sigma-Aldrich (Cat. #T7660) |
| FK866 hydrochloride hydrate | Sigma-Aldrich (Cat. #F8557) |
| β-Nicotinamide mononucleotide (NMN) | Sigma-Aldrich (Cat. #N3501) |
| 3x FLAG peptide lyophilized powder | Sigma-Aldrich (Cat. #F4799) |
| Ficoll PM400 | GE Healthcare (Cat. #17-0310-10) |
| Polyethylene glycol 4000 | Kanto Kagaku (Cat. #32828-02) |
| Poly(ADP-ribose) (PAR) Polymer | Trevigen (Cat. #4336-100-01) |
| β-Nicotinamide adenine dinucleotide (NAD) hydrate | Sigma-Aldrich (Cat. #N7004) |
| Phosphatase inhibitor cocktail I | This paper |
| Phosphatase inhibitor cocktail II | (Watanabe et al., 2018) |
| Nicotinamide | Sigma-Aldrich (Cat. #N0078) |
| Gallotannin | Sigma-Aldrich (Cat. #403040) |
| Hoechst 33258 | Dojindo (Cat. #343-07961) |
| Protein A conjugated to 10-nm gold particles | Cell Microscopy Center, University Medical Center Utrecht |
| Cell lines | |
| Human: Tetracycline-inducible Venus-ASK3-stably expressing HEK293A cells | This paper |
| Human: Tetracycline-inducible FLAG-ASK3-stably-expressing HEK293A cells | (Watanabe et al., 2018) |
| Human: HEK293A cells | Invitrogen |
| Oligonucleotides | |

| | |
|---|---|
| Control siRNA (Stealth RNAi Negative Control Medium GC Duplex #2) | Invitrogen (Cat. #12935-112) |
| NAMPT siRNA #1 (Stealth RNAi siRNA, target sequence: 5′-CCACCGACUCCUACAAGGUUACUCA-3′) | Invitrogen (Cat. #10620312) |
| NAMPT siRNA #2 (Stealth RNAi siRNA, target sequence: 5′-GAUCUUCUCCAUACUGUCUUCAAGA-3′) | Invitrogen (Cat. #10620312) |
| NAMPT siRNA #3 (Stealth RNAi siRNA, target sequence: 5′-GAAUAUUGAACUGGAAGCAGCACAU-3′) | Invitrogen (Cat. #10620312) |
| Expression plasmids | |
| pcDNA4/TO Venus-ASK3 (CDS of NM_001001671.3 with c.147C>T, c.574G>A) | This paper |
| pcDNA6/TR | Invitrogen (Cat. #V102520) |
| pcDNA4/TO EGFP-FLAG-ASK3 (CDS of NM_001001671.3 with c.147C>T, c.574G>A) | This paper |
| pcDNA3 ASK3-tdTomato (CDS of NM_001001671.3 with c.147C>T, c.574G>A) | This paper |
| pcDNA4/TO EGFP-FLAG | This paper |
| pcDNA4/TO EGFP-FLAG-ASK3(ΔN) (CDS of NM 001001671.3 with c.1 1,866del) | This paper |
| pcDNA4/TO EGFP-FLAG-ASK3(ΔC) (CDS of NM_001001671.3 with c.147C>T, c.574G>A, c.2,734_3,939del) | This paper |
| pcDNA4/TO EGFP-FLAG-ASK3(NT) (CDS of NM_001001671.3 with c.147C>T, c.574G>A, c.1,867_3,939del) | This paper |
| pcDNA4/TO EGFP-FLAG-ASK3(KD) (CDS of NM_001001671.3 with c.1_1,866del, c.2,734_3,939del) | This paper |
| pcDNA4/TO EGFP-FLAG-ASK3(CT) (CDS of NM_001001671.3 with c.1_2,712del) | This paper |
| pcDNA4/TO EGFP-FLAG-ASK3(ΔCCC) (CDS of NM_001001671.3 with c.147C>T, c.574G>A, c.3,535_3,675del) | This paper |
| pcDNA4/TO EGFP-FLAG-ASK3(ΔCLCR) (CDS of NM_001001671.3 with c.147C>T, c.574G>A, c.3,838_3,879del) | This paper |
| pcDNA4/TO EGFP-FLAG-ASK3(ΔCCCΔCLCR) (CDS of NM_001001671.3 with c.147C>T, c.574G>A, c.3,535 3,675del, c.3,838 3,879del) | This paper |
| pcDNA3/GW ANKRD52-Venus (CDS of NM_173595.3) | This paper |
| pcDNA4/TO FLAG-ASK3 (CDS of NM_001001671.3 with c.147C>T, c.574G>A) | (Watanabe et al., 2018) |
| pcDNA4/TO EGFP-ASK3 (CDS of NM_001001671.3 with c.147C>T, c.574G>A) | (Watanabe et al., 2018) |
| pcDNA3/GW FLAG-NAMPT (CDS of NM_005746.2) | This paper |
| pcDNA3/GW FLAG-NAMPT(S199D) (CDS of NM_005746.2 with c.595TC>GA) | This paper |
| pcDNA3/GW FLAG-NAMPT(S200D) (CDS of NM_005746.2 with c.598TC>GA) | This paper |
| pcDNA3/GW YFP-PPP6R3 (CDS of NM_001164161.1) | (Watanabe et al., 2018) |
| pcDNA3/GW PPP6C-HA (CDS of NM_002721.4) | (Watanabe et al., 2018) |
| pcDNA3/GW FLAG-ASK3 (CDS of NM_001001671.3 with c.147C>T, c.574G>A) | (Naguro et al., 2012) |
| pcDNA3/GW PARG-HA (CDS of NM_001303486.1) | This paper |
| pcDNA3/GW PARG(E673A/E674A)-HA (CDS of NM_001303486.1 with c.2,018AAGAA>CCGCC) | This paper |
| pcDNA3/GW PARG-Venus (CDS of NM_001303486.1) | This paper |
| pcDNA3/GW PARG(E673A/E674A)-Venus (CDS of NM_001303486.1 with c.2,018AAGAA>CCGCC) | This paper |
| pcDNA3/GW HA-PABPC1 (CDS of NM_002568.4) | This paper |
| pcDNA3/GW Venus-DCP1A (CDS of NM_018403.7) | This paper |
| pcDNA4/TO EGFP-FLAG-ASK3(CTΔIDR1) (CDS of NM_001001671.3 with c.1_2,712del, c.2,788_2,898del) | This paper |
| pcDNA4/TO EGFP-FLAG-ASK3(CTΔIDR2) (CDS of NM_001001671.3 with c.1_2,712del, c.3,427_3,540del) | This paper |
| pcDNA4/TO EGFP-FLAG-ASK3(CTΔIDRs) (CDS of NM_001001671.3 with c.1_2,712del, c.2,788_2,898del, c.3,427_3,540del) | This paper |
| pcDNA4/TO EGFP-FLAG-ASK3(CTΔCC) (CDS of NM_001001671.3 with c.1_2,712del, c.3,535_3,675del) | This paper |
| pcDNA4/TO EGFP-FLAG-ASK3(CTΔSAM) (CDS of NM_001001671.3 with c.1_2,712del, c.3,718_3,909del) | This paper |
| pcDNA4/TO EGFP-FLAG-ASK3(CTΔLCR) (CDS of NM_001001671.3 with c.1_2,712del, c.3,838_3,879del) | This paper |

| | |
|---|---|
| pcDNA4/TO EGFP-FLAG-ASK3(CTΔCCΔSAM) (CDS of NM_001001671.3 with c.1_2,712del, c.3,535_3,675del, c.3,718_3,909del) | This paper |
| pcDNA4/TO EGFP-FLAG-ASK3(CTΔCCΔLCR) (CDS of NM_001001671.3 with c.1 2,712del, c.3,535 3,675del, c.3,838 3,879del) | This paper |
| pcDNA3/GW CD38-HA (CDS of NM_001775.3) | This paper |
| pcDNA3/GW SIRT2-HA (CDS of NM_030593.2) | This paper |
| pcDNA3/GW HA-PARP1 (CDS of NM_001618.3) | This paper |
| pcDNA4/TO EGFP-FLAG-ASK3(PBM1; R58A/R59A) (CDS of NM_001001671.3 with c.147C>T, c.172CGGCGG>GCCGCC, c.574G>A) | This paper |
| pcDNA4/TO EGFP-FLAG-ASK3(PBM2; R203A/R204A) (CDS of NM_001001671.3 with c.147C>T, c.574G>A, c.607AGACGA>GCCGCC) | This paper |
| pcDNA4/TO EGFP-FLAG-ASK3(PBM3; R252A/K253A/R255A) (CDS of NM_001001671.3 with c.147C>T, c.574G>A, c.754CGGAAA>GCCGCC, c.763AGA>GCC) | This paper |
| pcDNA4/TO EGFP-FLAG-ASK3(PBM4; R332A/R333A) (CDS of NM_001001671.3 with c.147C>T, c.574G>A, c.994AGGAGA>GCCGCC) | This paper |
| pcDNA4/TO EGFP-FLAG-ASK3(PBM5; R391A/K392A) (CDS of NM_001001671.3 with c.147C>T, c.574G>A, c.1,171CGCAAA>GCCGCC) | This paper |
| pcDNA4/TO EGFP-FLAG-ASK3(PBM6; R424A/K425A) (CDS of NM_001001671.3 with c.147C>T, c.574G>A, c.1,270AGGAAA>GCCGCC) | This paper |
| pcDNA4/TO EGFP-FLAG-ASK3(PBM7; R436A/K437A) (CDS of NM_001001671.3 with c.147C>T, c.574G>A, c.1,306AGAAAA>GCCGCC) | This paper |
| pcDNA4/TO EGFP-FLAG-ASK3(PBM8; R493A/R494A/K496A/K497A) (CDS of NM_001001671.3 with c.147C>T, c.574G>A, c.1,477CGGCG>GCCGC, c.1,686AAGAAA>GCCGCC) | This paper |
| pcDNA4/TO EGFP-FLAG-ASK3(PBM9; K797A/R798A) (CDS of NM_001001671.3 with c.147C>T, c.574G>A, c.2,389AAACGT>GCCGCC) | This paper |
| pcDNA4/TO EGFP-FLAG-ASK3(PBM10; K895A/R896A) (CDS of NM_001001671.3 with c.147C>T, c.574G>A, c.2,683AAACGT>GCCGCC) | This paper |
| pcDNA3/GW HA-ASK3 (CDS of NM_001001671.3 with c.147C>T, c.574G>A) | (Naguro et al., 2012) |
| pcDNA3/GW HA-ASK3(PBM4; R332A/R333A) (CDS of NM_001001671.3 with c.147C>T, c.574G>A, c.994AGGAGA>GCCGCC) | This paper |
| pcDNA4/TO EGFP-FLAG-WWE (c.247–549 in CDS of NM_030963.3) | This paper |
| pcDNA3 ASK3(PBM4; R332A/R333A)-tdTomato (CDS of NM_001001671.3 with c.147C>T, c.574G>A, c.994AGGAGA>GCCGCC) | This paper |
| pcDNA3 ASK3(CT)-tdTomato (CDS of NM_001001671.3 with c.1_2,712del) | This paper |
| Software | |
| IUPred2A | (Mészáros et al., 2018) (URL https://iupred2a.elte.hu/) |
| Block-iT RNAi Designer | Invitrogen (current URL https://rnaidesigner.thermofisher.com/rnaiexpress/) |
| GNU Image Manipulation Program (GIMP; ver. 2.8.22) | GIMP Development Team (URL https://www.gimp.org/) |
| Fiji/ImageJ (ver. 2.0.0) | (Schindelin et al., 2012) (URL https://fiji.sc/) |
| Fiji plugin: TrackMate (ver. 4.0.1) | (Tinevez et al., 2017) (URL https://imagej.net/TrackMate) |
| RStudio (ver. 1.2.1335) | RStudio (URL https://www.rstudio.com/) |
| Language: R (ver. 3.6.0) | R Foundation (URL https://www.r-project.org/) |
| R package: tidyverse (ver. 1.2.1) | H. Wickham (RStudio team), URL https://www.tidyverse.org/) |
| R package: multcomp (ver. 1.4.10) | T. Hothorn *et al*. (URL http://multcomp.R-forge.R-project.org) |
| R package: lawstat (ver. 3.3) | J. L. Gastwirth *et al*. (URL https://CRAN.R-project.org/package=lawstat) |
| R package: ggpubr (ver. 0.2.4) | A. Kassambara (URL https://rpkgs.datanovia.com/ggpubr/) |
| R package: RColorBrewer (ver. 1.1.2) | E. Neuwirth (URL https://CRAN.R-project.org/package=RColorBrewer) |
| R package: lemon (ver. 0.4.3) | S. M. Edwards (URL https://github.com/stefanedwards/lemon) |
| ExPASy ProtParam tool | (Gasteiger et al., 2005) (URL https://web.expasy.org/protparam/) |

| | |
|---|---|
| Language: Python (ver. 3.6.5) | Python Software Foundation (URL https://www.python.org/) |
| Python library: NumPy (ver. 1.14.5) | NumPy Developers (URL https://numpy.org/) |
| Python library: pandas (ver. 0.23.3) | PyData Development Team (URL https://pandas.pydata.org/) |
| Python library: Pillow (ver. 5.2.0) | A. Clark *et al*. (URL https://python-pillow.org/) |
| Python library: Matplotlib (ver. 3.0.2) | J. Hunter *et al*. (URL https://matplotlib.org/) |
| Python library: seaborn (ver. 0.9.0) | M. Waskom (URL https://seaborn.pydata.org/) |
| Others | |
| 35 mmφ glass bottom dish | Matsunami (Cat. #D11130H) |
| 15 mmφ cover slip | Matsunami (Cat. #C015001) |
| 15x24 mm cover slip | Matsunami (Cat. #C01824) |

**Table S2. Statistical analysis**

Statistical tests, the number of samples and the sample sizes are indicated.

| Figure | Data | Number of Samples | Sample Size, *n* | Statistical Test |
|---|---|---|---|---|
| Figure 4 | | | | |
| 4F | FRAP rate at 180 sec | 2 (DMSO, FK866) | DMSO: 9, FK866: 8 (pooled from 3 independent experiments) | Brunner–Munzel's tests |
| 4F | FRAP rate at 180 sec | 3 (Control, + PARG WT, + PARG EA) | Control: 14, + PARG WT: 15, + PARG EA: 15 (pooled from 5 independent experiments) | Brunner–Munzel's tests with the Bonferroni correction |
| 4H | Normalized amount of condensate | 4 (Control, + 1.25 µM PAR, + 2.5 µM PAR, + 2.5 µM NAD) | 4 independent experiments | Dunnett's test |
| Figure S3 | | | | |
| S3A | Relative ASK3 activity of WT | 2 (Iso, Hyper) | 4 independent experiments | Paired *t*-test with the Bonferroni correction |
| S3A | Relative ASK3 activity of ΔC | 2 (Iso, Hyper) | 4 independent experiments | Paired *t*-test with the Bonferroni correction |
| S3A | Relative ASK3 activity of ΔCCC | 2 (Iso, Hyper) | 4 independent experiments | Paired *t*-test with the Bonferroni correction |
| S3A | Relative ASK3 activity of ΔCLCR | 2 (Iso, Hyper) | 4 independent experiments | Paired *t*-test with the Bonferroni correction |
| S3A | Relative ASK3 activity of ΔCCCΔCLCR | 2 (Iso, Hyper) | 4 independent experiments | Paired *t*-test with the Bonferroni correction |
| S3B | Relative ASK3 activity under hyperosmotic stress | 5 (siRNA (−), Control siRNA, NAMPT siRNA#1, NAMPT siRNA#2, NAMPT siRNA#3) | 5 independent experiments | Dunnett's test |
| S3C | Relative ASK3 activity under hyperosmotic stress | 4 (siRNA (−), Control siRNA, NAMPT siRNA#2, NAMPT siRNA#3) | 4 independent experiments | Dunnett's test |
| S3C | Relative OSR1 activity under hyperosmotic stress | 4 (siRNA (−), Control siRNA, NAMPT siRNA#2, NAMPT siRNA#3) | 4 independent experiments | Dunnett's test |
| S3D | Relative ASK3 activity under hyperosmotic stress | 8 (Control, NAMPT WT (+), NAMPT WT (++), NAMPT S199D (+), NAMPT S199D (++), NAMPT S200D (+), NAMPT S200D (++), Control (500 mOsm)) | 4 independent experiments | Dunnett's test |
| S3E | Relative ASK3 activity under hyperosmotic stress | 4 (Control, FK866, NMN, FK866 + NMN) | 5 independent experiments | Dunnett's test |
| S3F | Relative PPP6R3 interaction with ASK3 under hyperosmotic stress | 2 (DMSO, FK866) | 4 independent experiments | Paired *t*-test |
| S3F | Relative PPP6C interaction with ASK3 under hyperosmotic stress | 2 (DMSO, FK866) | 4 independent experiments | Paired *t*-test |
| S3F | Relative ASK3 activity under hyperosmotic stress | 2 (DMSO, FK866) | 4 independent experiments | Paired *t*-test |
| S3G | Relative PPP6R3 interaction with ASK3 under hyperosmotic stress | 2 (ultrapure water, NMN) | 4 independent experiments | Paired *t*-test |
| S3G | Relative PPP6C interaction with ASK3 under hyperosmotic stress | 2 (ultrapure water, NMN) | 4 independent experiments | Paired *t*-test |
| S3G | Relative ASK3 activity under hyperosmotic stress | 2 (ultrapure water, NMN) | 4 independent experiments | Paired *t*-test |
| S3H | Relative amount of PARsylated proteins | 2 (Control, NAMPT overexpression) | 3 independent experiments | Unpaired two-sample *t*-test with the Bonferroni correction |
| S3H | Relative amount of PARsylated proteins | 2 (DMSO, FK866) | 3 independent experiments | Unpaired two-sample *t*-test with the Bonferroni correction |

| | | | | |
|---|---|---|---|---|
| S3I | Relative ASK3 activity under hyperosmotic stress | 3 (Control, + PARG WT, + PARG EA) | 5 independent experiments | Dunnett's test |
| Figure S4 | | | | |
| S4A | Relative ASK3 activity under hyperosmotic stress | 2 (Control, + CD38) | 3 independent experiments | Unpaired two-sample *t*-test |
| S4B | Relative ASK3 activity under hyperosmotic stress | 2 (Control, + SIRT2) | 3 independent experiments | Unpaired two-sample *t*-test |
| S4C | Relative ASK3 activity under hyperosmotic stress | 2 (Control, + PARP1) | 4 independent experiments | Unpaired two-sample *t*-test |
| Figure S5 | | | | |
| S5C | Relative ASK3 activity of WT | 2 (Iso, Hyper) | 6 independent experiments | Paired *t*-test with the Bonferroni correction |
| S5C | Relative ASK3 activity of PBM1 | 2 (Iso, Hyper) | 6 independent experiments | Paired *t*-test with the Bonferroni correction |
| S5C | Relative ASK3 activity of PBM2 | 2 (Iso, Hyper) | 6 independent experiments | Paired *t*-test with the Bonferroni correction |
| S5C | Relative ASK3 activity of PBM3 | 2 (Iso, Hyper) | 6 independent experiments | Paired *t*-test with the Bonferroni correction |
| S5C | Relative ASK3 activity of PBM4 | 2 (Iso, Hyper) | 6 independent experiments | Paired *t*-test with the Bonferroni correction |
| S5C | Relative ASK3 activity of PBM5 | 2 (Iso, Hyper) | 6 independent experiments | Paired *t*-test with the Bonferroni correction |
| S5C | Relative ASK3 activity of PBM6 | 2 (Iso, Hyper) | 6 independent experiments | Paired *t*-test with the Bonferroni correction |
| S5C | Relative ASK3 activity of PBM7 | 2 (Iso, Hyper) | 6 independent experiments | Paired *t*-test with the Bonferroni correction |
| S5C | Relative ASK3 activity of PBM8 | 2 (Iso, Hyper) | 6 independent experiments | Paired *t*-test with the Bonferroni correction |
| S5C | Relative ASK3 activity of PBM9 | 2 (Iso, Hyper) | 6 independent experiments | Paired *t*-test with the Bonferroni correction |
| S5C | Relative ASK3 activity of PBM10 | 2 (Iso, Hyper) | 6 independent experiments | Paired *t*-test with the Bonferroni correction |
| S5D | Relative pulldowned ASK3 under hyperosmotic stress | 4 (WT-DMSO, WT-FK866, PBM4-DMSO, PBM4-FK866) | 4 independent experiments | Dunnett's test |
| S5D | Relative ASK3 activity under hyperosmotic stress | 2 (WT-DMSO, WT-FK866) | 4 independent experiments | Unpaired two-sample *t*-test with the Bonferroni corection |
| S5D | Relative ASK3 activity under hyperosmotic stress | 2 (PBM4-DMSO, PBM4-FK866) | 4 independent experiments | Unpaired two-sample *t*-test with the Bonferroni corection |
| S5F | FRAP rate at 180 sec | 3 (WT, PBM4, CT) | WT: 11, PBM4: 12, CT: 9 (pooled from 4 independent experiments) | Brunner–Munzel's tests with the Bonferroni correction |
| S5H | Relative amount of condensates | 2 (Control, + PAR) | 4 independent experiments | Unpaired two-sample *t*-test |
| S5H | Relative amount of condensates | 2 (Control, + PAR) | 4 independent experiments | Unpaired two-sample *t*-test |
| S5H | Relative amount of condensates | 2 (Control, + PAR) | 4 independent experiments | Unpaired two-sample *t*-test |

### Captions for Movies S1 to S4

**Movie S1.**

A computational simulation for the relationship between the grid space and the number/size of ASK3 clusters. Each frame was represented at every 10,000 iteration steps. Red squares: ASK3 units, black squares: obstacles.

**Movie S2.**

Dynamics and fusion of ASK3 condensates in Venus-ASK3-stably expressing HEK293A cells. After 5 min, the cells were exposed to hyperosmotic stress (500 mOsm). White bar: 20 μm.

**Movie S3.**

A computational prediction for ASK3 cluster disassembly after the grid space expansion. Each frame was represented at every 100,000 iteration steps. Red squares: ASK3 units, black squares: obstacles.

**Movie S4.**

Relationship between ANKRD52 and ASK3 condensates in HEK293A cells. After 5 min, the cells were exposed to hyperosmotic stress (500 mOsm). Magenta: ASK3-tdTomato, green: ANKRD52-Venus, white bar: 20 μm.
